# Ancient electrogenic survival metabolism of *β -proteobacterial* ammonia oxidizers for oxygen deficiency

**DOI:** 10.1101/2022.06.21.496957

**Authors:** Arda Gülay, Greg Fournier, Barth F. Smets, Peter R. Girguis

## Abstract

Oxygen availability is critical for microbes as some are obligatorily dependent on oxygen for energy conservation. However, aerobic microbes that live in environments with varying oxygen concentrations experience pressures over evolutionary time, selecting alternative energy metabolisms that relax the dependence on oxygen. One such capacity is extracellular electron transfer (or EET), which is the ability to transfer electrons from central metabolism to extracellular oxidants such as iron and manganese oxides. We posit that the *β-*proteobacterial ammonia-oxidizing bacteria, highly specialized lineages heretofore recognized as strict aerobes, can be capable of EET as they have been constantly observed in oxygen-limiting and depleted environments. Here, we show that a strictly aerobic ammonia-oxidizer, *Nitrosomonas communis*, utilized a poised electrode to maintain metabolic activity in anoxic conditions. The presence and activity of multi-heme cytochromes suggested that direct electron transfer is the mechanism underlying EET. Molecular clock models suggest that the ancestors of *β-*proteobacterial ammonia oxidizers appeared after the oxygenation of Earth when the oxygen levels were >10^-4^ *p*O_2_ (PAL), suggesting their aerobic origins. Phylogenetic reconciliations of gene and species trees show that the multi-heme c-type EET proteins in *Nitrosomonas* and *Nitrosospira* were acquired by gene transfer from *β-*proteobacteria during oxygen scarcity. The preservation of EET metabolism over billions of years under fluctuating oxygen levels and aspects of EET physiology in *β-*proteobacterial ammonia oxidizers might explain how they have been coped with oxygen stress and survived under oxygen deprivation.

**SIGNIFICANCE:** Metabolic versatility can permit typically aerobic microbes to survive in anaerobic conditions when oxygen is deficient as a terminal electron acceptor. This article demonstrates a previously unidentified anaerobic extracellular electron transfer metabolism that operates in aerobic *β*–proteobacterial ammonia oxidizers and reconstructs the evolutionary history of this metabolism, linking it to the early history of Earth’s oxygenation. Our approach contributes to the understanding of metabolisms in the N-cycle and their evolution on Earth, as well as how aerobic microbes manage to retain energy generation under oxygen-limiting or depleted conditions.

**AUTHOR CONTRIBUTIONS:** AG designed the physiological research with PRG and the phylogenetic research with GF; AG performed the research, AG analyzed the data with PRG and GF. AG wrote the paper, and all authors edited and approved the manuscript.

## INTRODUCTION

Energy conservation is essential to microbes for survival regardless of their type of metabolism (1). Microbes often employ diverse energy metabolisms to ensure their existence across a range of conditions, including non-optimal or extreme environmental conditions (2–4). The ability to transfer electrons from central metabolism to exogenous electron acceptor is called extracellular electron transfer, or EET (5). EET is enabled through a diversity of mechanisms, including redox-active soluble compounds(5) as well as multiheme c-type cytochromes (MHC), which are anchor proteins (6, 7) that link intracellular energy pathways to redox transformations of extracellular metal ions (5) or substrates –such as humic acids (8), soluble metal ion (9), dimethyl sulfoxide (10), as well as poised electrodes (11).

Transitions in molecular oxygen (O_2_) concentrations over Earth’s history have been a major force shaping modern microbial metabolisms (12). As the availability of O_2_ enabled the replacement of many enzymatic reactions central to anoxic microbial metabolism with aerobic respiratory chains (13), many lineages have evolved to be obligate aerobes. For example, - γ- and *β- proteobacterial* taxa that oxidize ammonia (NH_3_^+^) to nitrite (NO_2_^-^) via O_2_ reduction are considered to be strict aerobes (14). However, aerobic ammonia oxidizers are consistently observed in oxygen-limiting and depleted environments such as anoxic marine zones (15) and sediments (16, 17), as well as engineered systems (18). Although a wide range of oxygen affinities for aerobic ammonia-oxidizing bacteria have been reported (19), how they are able to persist and survive under reductive stress during oxygen deficiency remains enigmatic.

Ammonia and methane-oxidizing bacteria are recognized to be evolutionarily related based on their bioenergetic metabolism proteins, such as copper membrane monooxygenases (20). Few modern anaerobic methane-oxidizing clades (ANME) are capable of performing direct electron transfer between species using MHCs (6). The deeply rooted phylogeny of archaeal ANME clades in time (21, 22) suggests MHCs may have a long evolutionary history before the Proterozoic eon.

A few studies have reported anaerobic reactive-N transformation processes in ammonia-oxidizing bacteria, mostly suggesting an alternative reductive metabolism such as nitrite reduction (23–26), but such metabolisms were linked to re-oxidation of NAD(P)H instead of any exogenous energy-yielding reaction (27). In an anoxic mixed culture-based electrochemical system, it has been claimed that *Nitrosomonas* spp. donates electrons to an insoluble electrode (28). If aerobic ammonia-oxidizing bacteria can donate extracellular electrons to solid electrodes, they have the potential to generate energy via EET metabolic couplings in nature in the absence of O_2_. We hypothesize that ammonia-oxidizing bacteria can switch from aerobic metabolism to anaerobic EET metabolism under anoxic conditions.

To test this hypothesis, we examined the electrochemical response of model ammonia-oxidizing bacteria under anoxic conditions. Cyclic voltammetry and chronoamperometry were applied to determine electrochemical ability and the redox behavior of ammonia oxidizers, and cytochrome reactive heme staining and metatranscriptomics were employed to identify the presence and activity of their MHCs. We applied molecular clock models, integrated with reconciliations of genes associated with EET, to species tree of taxa carrying multi-heme EET cytochromes to estimate the divergence times of ammonia-oxidizing bacteria clades and EET performing clades. These were calibrated with secondary constraints on archaeal and bacterial divergence times from recently published analyses. Our results show that i) *Nitrosomonas* transfers extracellular electrons to solid surfaces, ensuring its survival under oxygen deprivation, and ii) this capacity is ancient, likely tracing back to the appearance of *β*–proteobacteria 1172-1936 Ma and showing that ammonia oxidation likely persisted across oxic and anoxic niches across much of Earth history.

## Results and Discussion

### Multi-heme EET cytochromes are distributed in Nitrosomonas lineage

The mechanism of EET often relies on a connection of redox and structural proteins; some of those are well-characterized in a few model microbes (29). A network of proteins affiliated to periplasmic c-type cytochromes, integral outer-membrane β-barrel proteins, and outer-membrane-anchored c-type cytochromes were shown to facilitate EET in *Shewanella onediensis, Geobacter sulfurreducens, and Rhodopseudomonas palustris* (5). We searched the NCBI genome database for genes used in EET metabolism in ammonia-oxidizing bacteria to evaluate their potential EET capability by screening the homologs of model EET pathways (Supp. Table 1). Putative homologs to these known examples were queried using HMMER on NCBI and Ensembl Reference Sequence databases (30). No consistent homology was detected of such EET pathways in archaeal ammonia oxidizers. Detected homologs in ammonia-oxidizing bacteria were more similar in sequence to the enzymatic EET machinery of *Shewanella Onediensis* (31) than *Geobacter* and *Rhodopseudomonas* and were primarily identified in the *Nitrosomonas* genus (Supp. Fig.1). *Nitrosomonas* species comprising the complete EET machinery formed a single coherent phylogenetic clade in the species tree (Figure 1), suggesting that the putative EET metabolism of *Nitrosomonas* appears to be a shared ancestral trait of this group (32). The γ-proteobacterial ammonia oxidizer *Nitrosococcus Halophilus* was also identified as carring a full set of EET metabolism genes.

**Figure 1:**
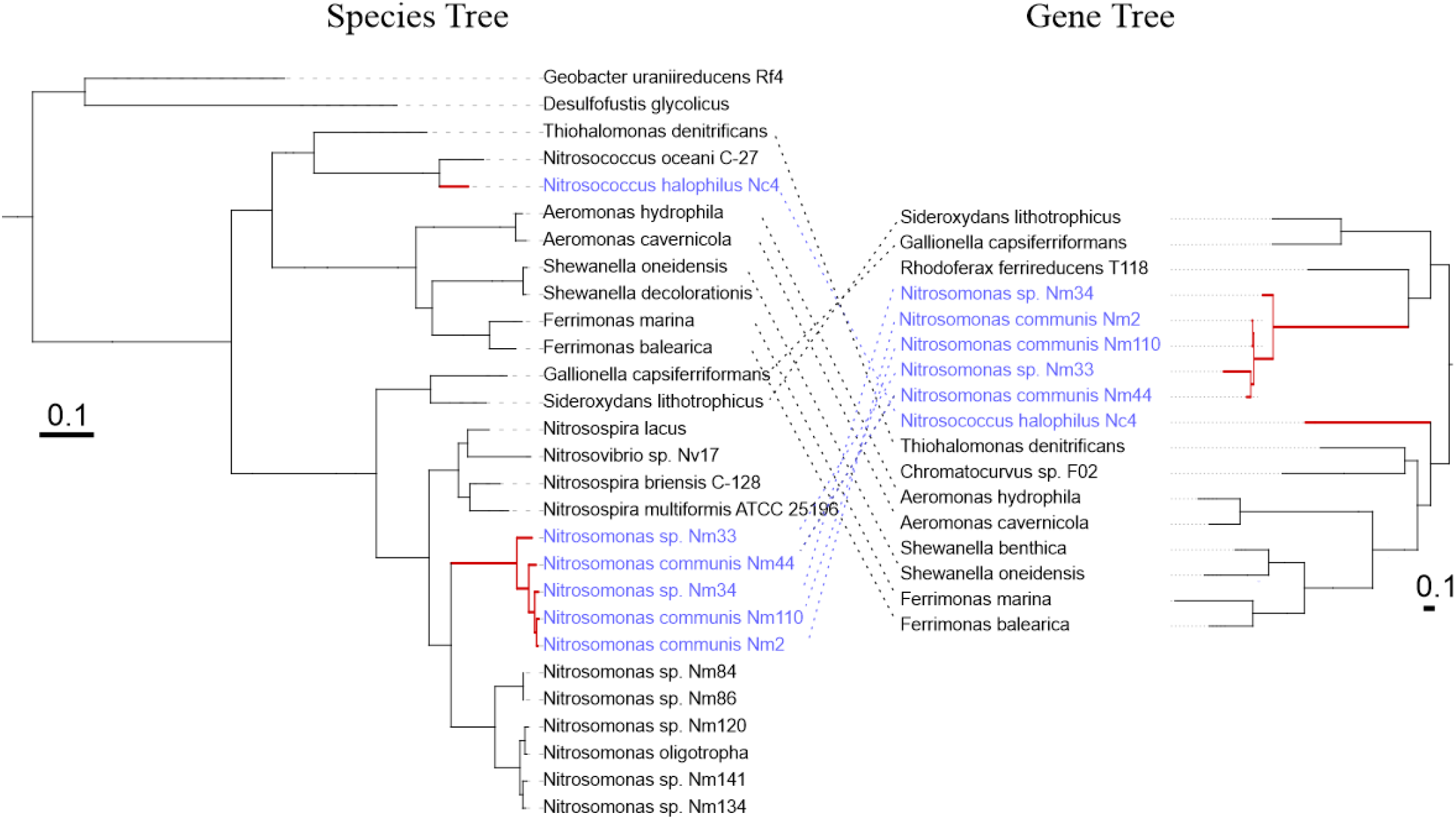
Comparison of phylogenies for species tree and EET metabolism gene tree. Species trees were generated from concatenated alignments of ribosomal protein sequences. EET gene trees were constructed from the concatenated alignments of proteins orthologous to identified EET components in ammonia-oxidizing bacteria, complete MTR pathway, including periplasmic c-type cytochromes (Q8EG35), integral outer-membrane β-barrel proteins (Q8EG34), and outer-membrane-anchored c-type cytochromes (Q8EG33). Ammonia oxidizers with (Blue) and without (Green) a known EET pathway.

We then examined multi-heme cytochromes (MHC) -with at least two “C-X-X-X-C-H” heme-binding motifsin ammonia-oxidizing bacteria (7) and compared the total number of MHC and heme-binding motifs in strains within the detected *Nitrosomonas* clade to other *Nitrosomonas strains* (Supp. Fig. 2). A total of 25 and 21 MHC carrying 87 and 70 heme-binding motifs were identified in *Nitrosomonas communis* Nm2 and *Nitrosococcus halophilus* Nc4, comprising homologs of the EET pathway, respectively, while other ammonia oxidizers, on average, contain 16 MHC with 57 heme-binding motifs (Supp. Table 2). The intense presence of MHCs with the presence of homologs of EET model metabolism among *Nitrosomonas* suggests that ammonia-oxidizing bacteria, *Nitrosomonas* and *Nitrosococcus*, can transfer electrons to surfaces under limited oxygen conditions. To test this hypothesis, we forwarded the live strains of *Nitrosomonas communis* Nm2 and *Nitrosococcus halophilus* Nc4 to physiological examination via bioelectrochemical incubations under oxygen-deficient conditions.

### Nitrosomonas donates electrons to solid electrodes in an anaerobic environment

We measured the electrochemical response of *N. communis* and *N. Halophilus* under anaerobic and ammonium supplemented conditions (Figure 2A). The anodic chronoamperometric (CA) current at 0.3V versus AgCl was stable at 4.7 ± 2 μA over ten days in electrochemical incubations of *N. communis*, while abiotic control consistently yielded 2.8 ± 0.7 μA over ten days of incubation (Figure 2B). These data suggest that the graphite felt electrode served as an electron acceptor for *N. communis* in the presence of ammonia, generating a constant anodic current at all live replicates higher than the controls. Notably, we did not observe an increase in the anodic current of live incubations, suggesting that the microbial electroactivity over this time course was insufficient to support rapid cell growth (33). *N. communis* may have longer doubling times while using an anaerobic energy metabolism, previously reported for *Nitrosomonas europaea* (6.5 days vs. 9 days in aerobic conditions; 34). The stable current could also be related to limited extracellular electron transfer by *N. communis* (35). *N. Halophilus* did not produce currents higher than control incubations (Supp. Fig. 3), indicating that *N. Halophilus* cannot perform EET at given conditions.

Electrochemical characteristics of *N. communis* were further analyzed by cyclic voltammetry (CV) via 1 mV/s scanning rate (Fig. 2C), specifically to determine the redox-active compounds of *N. communis*. Anodic CV peaks that appeared at the control incubations were consistently lower than at the live incubations, through which *N. communis* enabled more efficient electron transfer to the anode than the control (36). Interestingly, an anodic CV peak was observed in live incubations at the potential of 0.25V, close to the applied potential in the electrochemical experiments (Fig.2D). As all other anodic redox peaks were observed in liveand control incubations together, the redox signal at 0.25V appeared to be affiliated with *N. communis*(37). Such redox response suggests that the mechanism responsible for the electrochemical activity of *N. communis* involved redox compounds or proteins that developed under O_2_ deficient conditions. Furthermore, the current generation and redox activity indicated that *N. communis* could activate its extracellular electron transfer metabolism; however, the consistently steady CA trend suggests that such metabolism did not support the significant cell growth.

**Figure 2.**
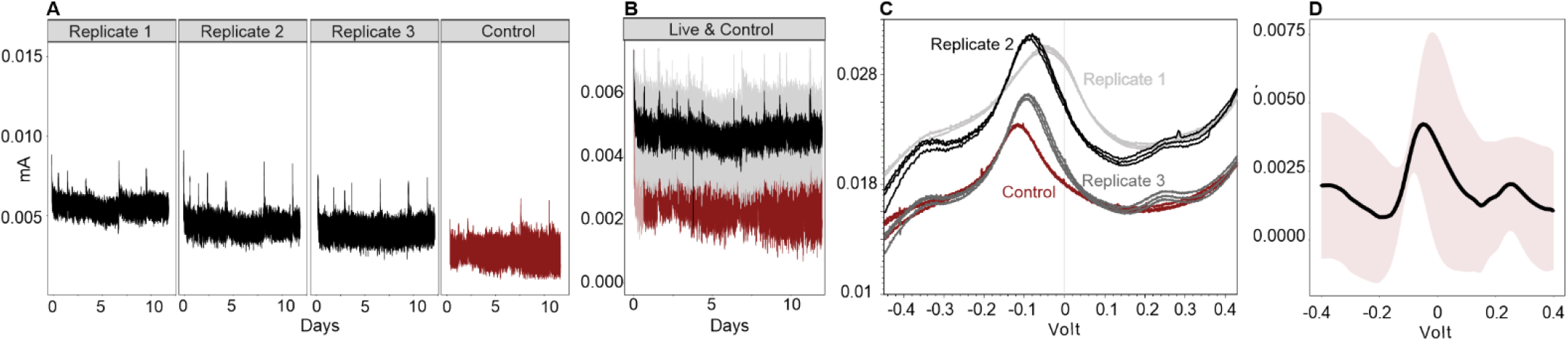
Electrochemical characteristics of N. communis on graphite felt electrode. (A) Anodic currents were measured using an electrode poised at 0.3V versus AgCl supplemented with 1 mM NH_4_^+^. (B) Mean anodic current from replicate runs of N.Communis (black) with standard deviation (grey) and abiotic control (red). (C) Cyclic voltammograms were measured at a scan rate of 1 mV s^-1^ ten days after initiating the incubation. (D) Mean cyclic voltammogram from replicates runs of N.Communis normalized by subtracting the abiotic control CV as a baseline. The smooth pink layer represents the standard deviation.

### Outer membrane and multi-heme cytochromes were active in anode respiring N. communis

Heme staining coupled with TEM imaging was applied to ascertain the presence and distribution of outer membrane heme-containing metalloprotein(s) cytochromes of electro-active *N. communis* after ten days of incubation without O_2_(7, 38). In cytochrome-specific 3,3′-diaminobenzidine (DAB)–H_2_O_2_ staining, heme metal-centers catalyze the formation of a DAB polymer which has a high binding affinity to OsO_4_(38). Thin sections of *N. communis* showed the apparent formation of heme-bound peroxidase, whereas *N. Halophilus*, which showed no electrochemical response, exhibited no heme-bound peroxidase activity (Figure 3A). This suggests that outer membrane cytochromes of *N. communis* were developed during the electrochemical incubation and potentially involved in the extracellular electron transfer mechanism (39). The control sections of *N. communis* without the amendment of DAB stain yielded no heme-bound peroxidase signal, indicating that DAB successfully reacted with heme centers of *N. communis* in non-control TEM sections. Outer membrane cytochromes are often an essential component of direct extracellular electron transfer between microbes and solid surfaces (29); the activity of outer membrane cytochromes further suggests that *N. communis* transferred its electrons to the solid electrode through direct electron transfer. The cytochrome activity of electrochemically incubated *N. communis* was further assessed via metatranscriptomics. Among 25 detected multi-heme cytochromes (≥2 heme motifs) in the genome of *N. communis* (Supp. Table 3), 19 were expressed after ten days of electrochemical incubation without oxygen, in line with electrochemical measurements and heme stain assays (Figure 3B). Furthermore, the transcript library of *N. communis* revealed the activity of genes affiliated with anaerobic metabolism; the highest expression levels were detected for P460 (Supp. Table 4), a cytochrome converting NH_2_OH to N_2_O (40), indicating its potential role in extracellular electron transfer activity. In line with the enzymatic mechanism of cytochrome p460 in *Nitrosomonas Europhea* (40), NH_2_OH oxidoreductase activity was also detected, suggesting p460 dependence on NH_2_OH oxidoreductase activity during EET. While a future isotopic tracing and a comprehensive transcriptomic effort may ultimately provide a complete understanding of the enzymatic EET pathway of *N. Communis*, our results suggested that cyc. P460 can be the enzymatic bridge between ammonia oxidation and the extracellular electron release of *N. communis*. In addition, the low expression of carbon fixation genes (Supp. Table 5) was consistent with the non-increasing anodic currents from CA results (Figure 2A-B), pointing out that the cell growth was not supported.

**Figure 3:**
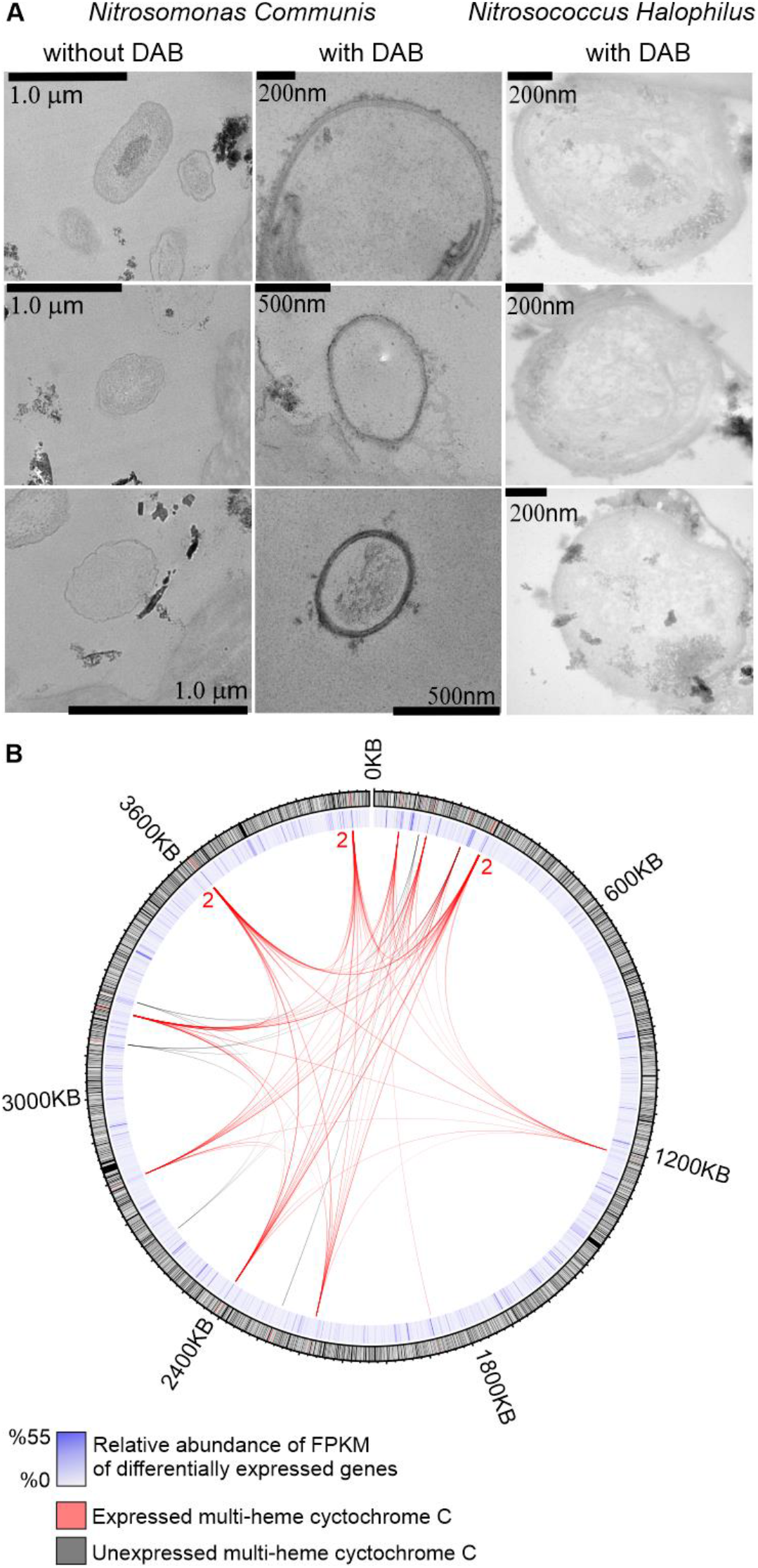
(Left) Transmission electron microscopy images of N. communis and N. halophilus cells stained with cytochrome-reactive DAB-H_2_O_2_ and without DAB. (Right) Expressed genes of N. communis and their expression levels (the inner circle) were shown with their genome position (the outer circle). Expressed (red) and unexpressed (black) multi-heme (>2) c-type cytochromes (MHC) were shown as net inside the circle.

### *β-proteobacterial* ammonia-oxidizing bacterial lineages are over 1.7 billion years old

While the physiological evidence indicated the active metabolism of EET in *Nitrosomonas* spp., we run phylogenomics to infer the importance of EET metabolism to ammonia-oxidizing bacteria by (i) dating the EET metabolism and ammonia oxidizers and (ii) reconstructing the evolutionary history of EET metabolism aligned with the evolution of O_2_ concentrations on Earth.We first applied molecular dating on a species tree (30, 41, 42), including *Nitrosomonas* spp., to track down the origination of ammonia-oxidizing bacterial clades and compare the timing of such events with other EET clades, such as *Shewanella*. Species tree was constructed with archaeal and bacterial taxa preserving EET metabolism and taxa involved in molecular clock calibration, affiliated to *Euryarcheota, Proteoarchaeota, Protobacteria, Cyanobacteria, Plantomycetes*, and *Verrucomicrobia*. The divergence times of the species tree were calibrated using secondary constraints for the age of the archaeal root (4.38−3.46 Gya) and the crown ages of methanogenic archaeal clades, including *Methanosarcinales, Methanococcoides, and Methanohalobium*, as published by Wolfe et al. 2018 (42). Younger bacterial clades such as *Aeromonas* (43) and *Vibrio* (44) were further used to calibrate the species tree together with *Nostocales*, using a fossil calibrated age estimate for this group (42, 45). We also applied molecular dating to the MtrA gene tree (Supp. Fig. 4) using the calibrations for *Aeromonas, Vibrio*, and clade 6 *Cyanobacteria* (46).

As evolutionary models often substantially impact the age estimates of molecular clocks (47), we used uncorrelated and autocorrelated branch-specific rate models, as well as uniform and birth-death priors on the relative age distributions of divergences to estimate the uncertainty of age estimates (48). Furthermore, a model assessment method was applied to determine the most compatible model with detected HGT events (Supp. Fig. 5) as described previously (48). CIR process model with a uniform prior resulted in the highest compatibility, where 94% of the posterior chronograms tested under CIR+UNI fulfilled HGT constraints (Supp. Table 5). Similarly, the CIR model with birth-death prior showed 93% compatibility, which is very close to the CIR+UNI; we, therefore, considered estimations that covered the time range of both models in our interpretations. The compatibility of other models ranged from 80% to 23%.

Molecular clock estimates under the CIR+UNI and CIR+BD models recover *Nitrosomonas* and *Nitrosospira* clades appearing during the Paleoproterozoic, between 2247 and 1556 Mya, while the *Nitrosococcus* group originated more recently, during the Neoproterozoic between 1558 and 426 Mya (Figure 4). The origination of *β* -proteobacterial ammonia oxidizers is consistent with the appearance of their EET metabolism, as indicated by the molecular dating analysis of the MtrA gene tree (Supp. Fig. 4). The *Shewanella* lineage in possession of EET metabolism was observed to originate during the Mesoproterozoic between 1686 and 1089 Mya, indicating that ammonia-oxidizing bacteria likely obtained EET metabolism before *Shewanella*. The appearance of *β*proteobacterial ammonia-oxidizing bacteria is after the Great Oxygenation Event (GOE), when the atmospheric oxygen level was more than >5-10% present atmospheric level (PAL) and before the atmospheric oxygen levels dropped again to 0.1% PAL (49–51) suggesting that the ancestor of such lineage was likely to be aerobic. These age estimates suggest that bacterial ammonia oxidizers acquired their metabolisms and diversified at different times during the Proterozoic, while the ancestral archaeal ammonia oxidizer of terrestrial *Nitrosocaldus, Nitrososphaera*, and *Nitrosocosmicus* lineages was estimated to be somewhat more ancient, and older than previously reported (2328-3100 Ma) (52).

### Ancestral ammonia-oxidizing bacteria acquired EET metabolism by HGT under the oxygen limitation

The early appearance of β proteobacterial ammonia-oxidizing bacteria raised the possibility that the functional EET metabolism detected in their modern state has been conserved for a long time. To date and construct a model of EET evolution, we compared the domain family tree of periplasmic c-type MHC (MtrA) to the dated species phylogeny using reconciliation, a technique that recognizes well-supported speciation, duplication, transfer, and loss events (53, 54) during the evolution of the EET MHC using ecceTERA v1.2.4 (35; Fig. 5). Among all the domain family trees of EET metabolism components, periplasmic MHCs were conserved and widely distributed among the tree of life (Supp. Fig. 1-6-7) and hence used to reconstruct the evolution of EET metabolism in the tree of prokaryotic life. Date estimates of key HGT recipient nodes (Supp. Fig. 8) were given under CIR, LN, and UGAM models with uniform and birth-date priors (Figure 5).

**Figure 4.**
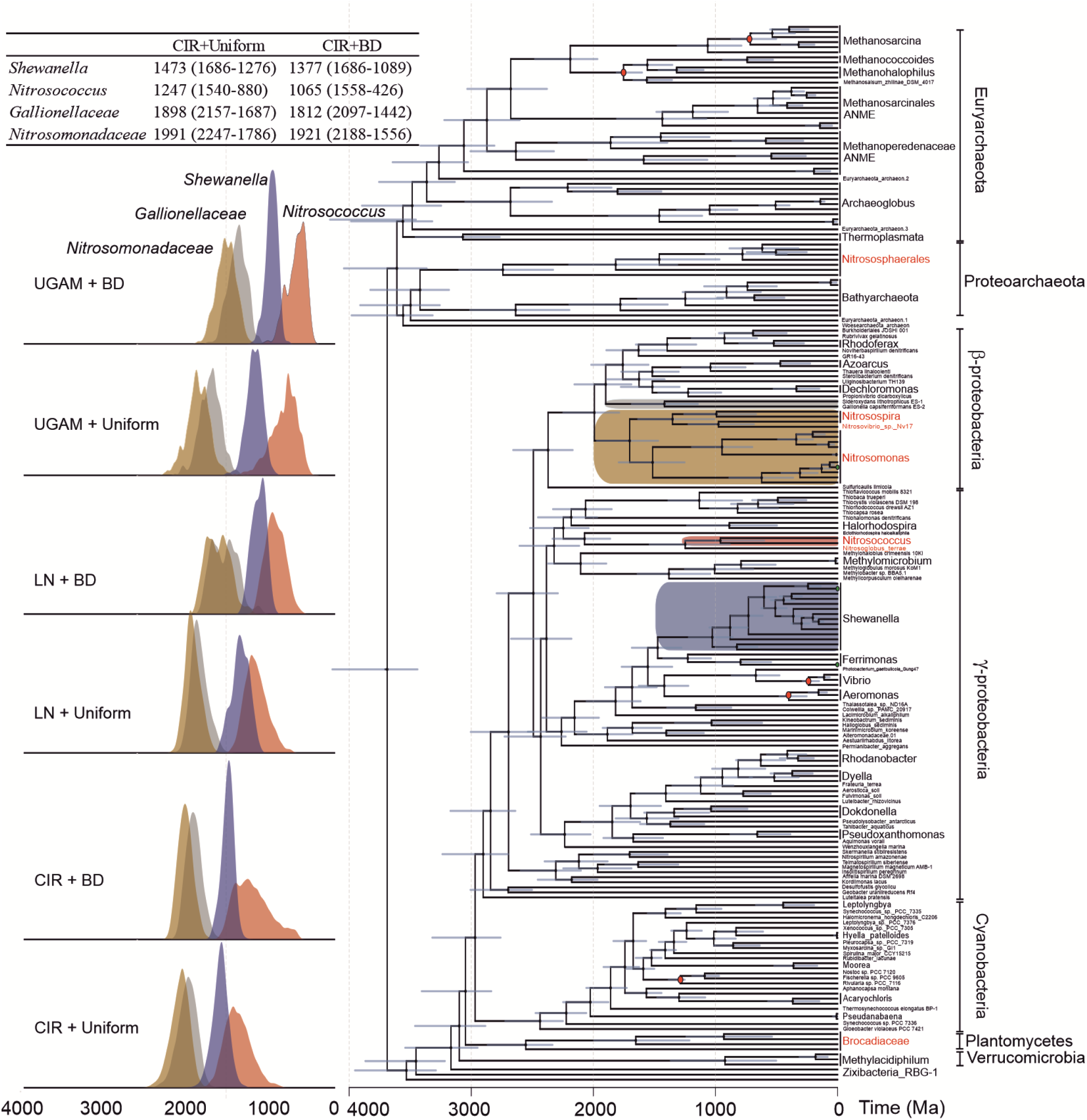
Time-calibrated species tree with ammonia-oxidizing clades highlighted. The depicted chronogram is for the CIR rate model with a uniform distribution, which was selected using the HGT compatibility method by Fournier et al. 2021. Key bacterial clades with putative EET metabolisms are highlighted. Calibrations are indicated by red-filled circles. Posterior distributions were generated by sampling the Markov chain Monte Carlo analysis every 1,000 generations, with a 25% burn-in. Blue bars show uncertainty (95% CI). The age distributions of key nodes for the timing of the diversification of Nitrosomonadaceae, Gallionellaceae, Shewanella, and Nitrosococcus lineages are provided on the left side graph under six different evolutionary models. The mean age estimates for a selection of key nodes under CIR+Uniform and CIR+BD models are provided in the table with 95% CI levels.

**Figure 5.**
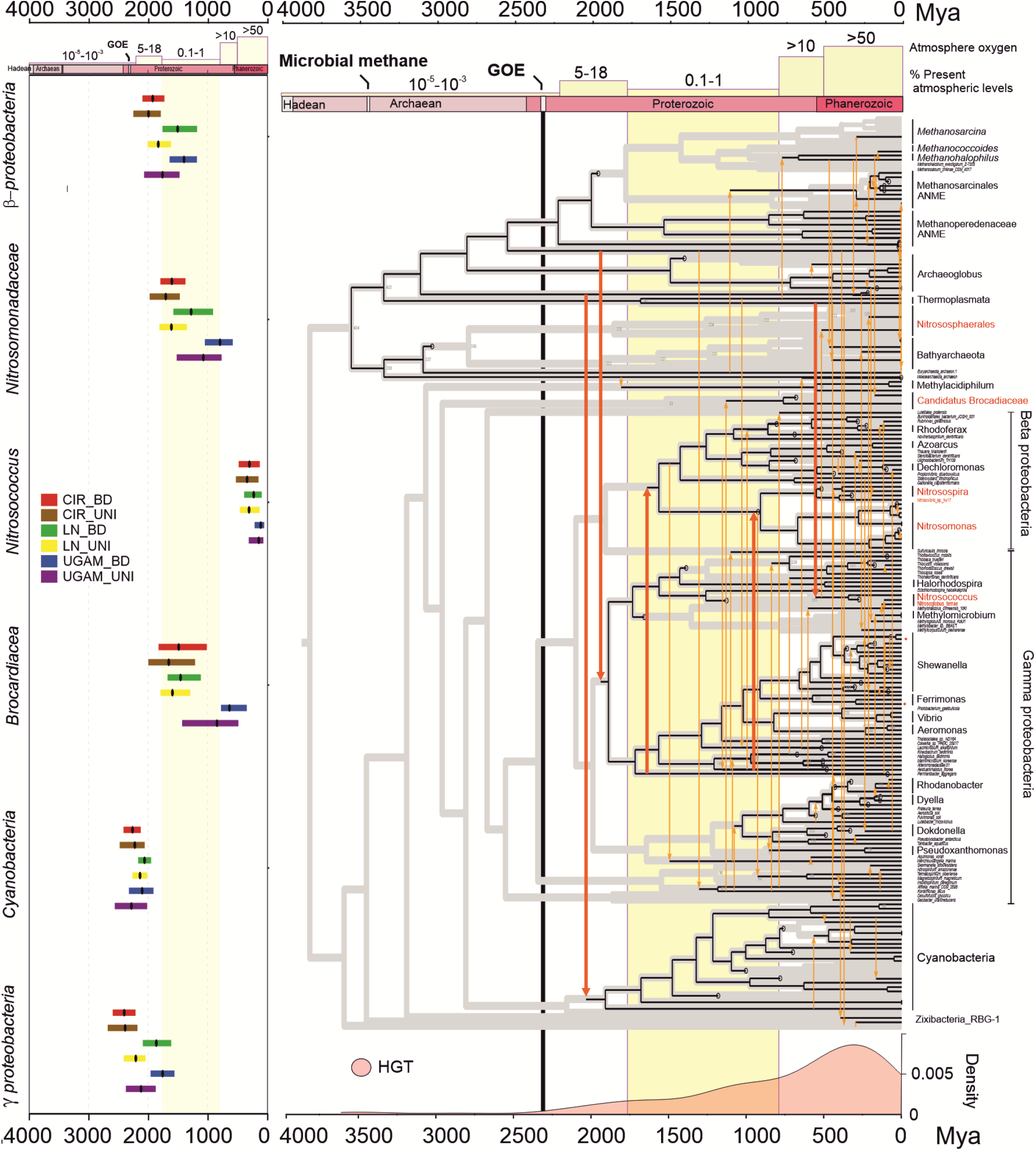
Visualized reconciliation of the electron transfer gene tree (MtrA) onto the dated species tree in Figure 5 constructed from concatenated ribosomal genes. Black lines represent vertical inheritance within the genomes of the species tree, whereas orange lines represent the gene transfer events. Red lines represent the important gene transfer events for bacterial clades and ammonia-oxidizing bacteria (Red species). Yellow boxes represent atmospheric oxygen levels over time. The bottom graph shows the frequency of gene transfer events of the electron transfer gene over time. Taxa are abbreviated as per Figure 5. A summary of the dates of the reconciled key nodes received the MtrA gene estimated with six evolutionary models was given on the left.

The reconciled roots for periplasmic MHC, MtrA, map to node 424 (3363-4045 Ma), the last common ancestor of Euryarchaeota and TACK in Figure 6. We further evaluated the EET root position by forcing a posterior root age for the last common ancestor of the MtrA (node 424) in the MtrA gene tree using the age estimates for the crown groups from the species tree as secondary calibrations (Supp. Fig. 9). The molecular clock applied to the MtrA gene tree yielded the posterior age estimate of 3348 – 4010 Ma to the last common MtrA ancestor in the species tree, which is consistent with the age estimate of the reconciled tree (3363- 4045 Ma). The origins of modern EET metabolism in *Archaeoglobus, Thermoplasmata*, and anaerobic methane-oxidizing archaea (ANME) affiliated to *Methanoperedenaceae* reached out to the earliest MtrA; the MHC-driven direct interspecies electron transfer (DIET) between ANME and sulfate-reducing Deltaproteobacteria (SRB) as well as the potential metal-reducing metabolism of ANME has been recently documented (6).

It is important to mention that the inferences of deep-rooted ancestry from phylogenetic reconciliation are sensitive to the sampled gene diversity in that the origins of MtrA may not have been covered by the observed species tree due to the shortage of identified genomes so far. The HGT events and recipients inferred by the reconciliations, on the other hand, were less prone to biases from rooting, reconciliation, and sampling. The reconciliation of the MtrA phylogeny to the species tree suggests that the distribution of MtrA genes in the bacterial tree of life can be explained by major gene transfer events between archaea to bacteria (Figure 5, red arrows). The earliest lateral gene transfers mapped from (1) *Thermoplasmata* to *Cyanobacteria* and (2) *Methanoperedenaceae to Gamma-proteobacteria*. Those possible HGT events occurred after GOE around 2000 Mya (Figure 5).

Phylogenetic reconciliation suggests that the shared ancestor of *Nitrosomonas* and *Nitrosospira* acquired the periplasmic EET MHC from *gamma-proteobacteria*, which is also the case for the *beta-proteobacteria* (Figure 5). The lateral gene transfer event to ammonia-oxidizing bacteria, *Nitrosomonadaceae* and Candidatus *Brocadiaceae*, consistently maps in all models into the ancestral node during oxygen limitation in all estimations when the surface oceans were largely anoxic due to the critical drop of atmospheric oxygen levels (47, 48). The oxygen-reducing ancestral *β-proteobacterial* ammonia oxidizer acquired the anaerobic EET metabolism under oxygen limitation and preserved it to now. As in *Methanoperedenaceae ANME*, our study indicated that the EET metabolism is still functional in *β-proteobacterial* ammonia-oxidizers today.

Gene loss is a major evolutionary process shaping bacterial genomes (56, 57); it often depends on the importance of the gene to the microbe (58–60), and it correlates with time (57). Environmental shifts mostly lead to gene loss events if a gene becomes non-adaptive due to the new environment where its host resides (61, 62). The persistent conservation of an anaerobic energy metabolism regardless of variant oxygen levels over eons suggests that EET metabolism is an essential metabolic component of *β-proteobacterial* ammonia oxidizers. The electrochemical behavior of EET metabolism and transcriptome of EET selective incubations in *Nitrosomonas* spp. suggested that the ammonia oxidizers activate EET under oxygen stress and can place its metabolism to a dormant state, such as minimizing the acquired energy for cell growth. Physiological and phylogenomics indicated that the anaerobic EET metabolism provides a survival strategy for aerobic ammonia oxidizers while exposed to oxygen-deficient conditions.

## Material and Methods

### Bacterial strains and culture conditions

The strains used for this study were *Nitrosomonas Communis Nm2* [acquired from Dr. Lisa Stein’s lab] and *Nitrosococcus Halophilus* [Japan Collection of Microorganisms (JCM), 30413]. *N. communis* was grown in oxic media containing (g per liter): (NH_4_)_2_SO_4_: 0.66; NaCl: 0,58; KH_2_PO_4_: 0,05; MgSO_4_.7H_2_O: 0,05; CaCl_2_.2H_2_O: 0,14; KCl: 0,07. The pH was adjusted to 7.6. *N. Halophilus* was cultivated with JCM 1056 media. Cultures were incubated at 28 using a rotary shaker.

### Electrochemical incubation and analysis

Two-chamber electrochemical reactors were constructed using 100 mL glass Schott bottle sealed with butyl stopper. The compartments were separated using Nafion™ tubing (PermaPure LLC, USA), and the anode was equipped with a custom-made AgCl reference electrode, and carbon felt (3 cm length, 3 cm width, 1.12 cm thickness, 0.72 m2/g surface area, Alfa-Aesar, USA) working electrode attached to a titanium wire (0.06 cm diameter; Sigma), whereas a platinum wire was used as the counter electrode at the cathode. Continuous gassing of N_2_/CO_2_ was provided to ensure the constant mixing, anaerobiosis, and pH, in reactors. Culture media were used as the electrolyte after replacing (NH_4_)_2_SO_4_ to NH_4_Cl and MgSO_4_.7H_2_O to MgCI_2_ and degassing with N_2_/CO_2_ for a day. *N*.*communis* and *N. Halophilus* were filtered (0.22 μM) from the aerobic media, washed, and resuspended in 5 mL anoxic electrolyte in an anaerobic chamber. Electrolyte solution then gently vortexed with carbon felt working electrode and left for 15 minutes of settling. The electrochemical reactor was then sealed with the culture-attached working electrode, remaining electrolyte, and all other components inside the anaerobic chamber and placed for the chronoamperometric run while purging N_2_/CO_2_.

For chronoamperometric measurements, VMP3 multi-channel potentiostat (BioLogic Company, France) was used on electrochemical reactors with an applied anode potential of 0.3V (vs. AgCl). Along with triplicate reactors of *N. Communis* and *N. Halophilus*, parallel control reactors were operated without biomass. Identical voltage settings were used for both inoculated and control reactors. At the end of the 12 days chronoamperometric run, Cyclic voltammetry (CV) was applied to characterize the redox behavior of the cultures. The scan range of CV was from 0.4 V to -0.4 V, with a scan rate of 1 mV/s, using the VMP3 multi-channel potentiostat. A low scan rate 1 mV/s was applied to minimize the background capacitive current and the kinetic limitations of interfacial electron transfer between the microbial cells and the electrode (63). The mean CV of *N. Communis* was calculated with QSoas (http://bip.cnrsmrs.fr/bip06/software.html) after subtracting the abiotic CV from all biotic CVs as a baseline.

### Cytochrome reactive staining and Transmission electron microscopy

*N. Communis and N. Halophilus* cells were fixed from 1×1×1 cm3 piece of working electrode at the end of 12 days of incubation using deaerated solutions containing 2% paraformaldehyde and 2.5% glutaraldehyde on ice. After fixation, cells were dislodged from the electrode material by gentle vortexing, and suspended cells were filtered through 0.22 μM Nuclepore™ Track-Etched polycarbonate filters. Filters were washed 5X with 1.5 mL 50 mM Na^+^-Hepes (pH 7.4, 35 g/liter NaCl). DAB-H_2_O_2_ and OsO_4_ staining were applied as described previously (6). After dehydration through a graded series of ethanol, DAB and OsO4 stained filters were placed first into 1:1 propylene oxide/ethanol solution and then into 100% propylene oxide solution. Epon812 was used as embedding resin for electron microscopy and polymerized at 60°C overnight with the filters after preincubation using 100 μL ethanol/Epon 812 (1:1) overnight.

### Transcriptomic library and analysis

Total RNA extraction was applied to 2×2×2 cm^3^ working electrode at the end of the 12 days incubation. The electrode material was immediately embedded into a “Trizol Reagent” (ThermoFisher Scientific) in FastPrep Tubes with Zirconia/Silica beads. Bead beating with MP Biomedicals™ FastPrep -24™ for 90s at 10 m/sec was then applied, followed by 200 μl chloroform addition to promote phase separation. With the clear phase separation and formation using centrifugation at 12,000 x g for 15 minutes at 4°C, the total RNA was extracted from the clear aqueous phase of the solution using Direct-zol™ RNA MiniPrep kit (Zymo Research) according to the manufacturer’s instructions. rRNA was removed with Ribo-Zero Plus kit (Illumina Inc.), and cDNA sequencing library was prepared using the TruSeq Stranded mRNA Library Prep Kit (Illumina), following the manufacturer’s guidelines. Sequencing was applied with NovaSeq 6000, S2 100PE generating ∼50M (25M per read) million 100bp pair-end reads per library.

Sequence quality control tool FastaQC (64) and sequence pre-processing tool Trimmomatic(65) were used to perform the quality check and process the raw paired-end reads. The published genome of *N. Communis* Nm2 (GenBank: GCA_001007935.1, RefSeq: GCF_001007935.1) was used as the reference genome and transcripts for this study. The alignments of processed pair-end reads were performed using HISAT2(66) and aligned sam files were then sorted and processed using SAMtools suit. The transcript counts were then performed using Stringtie2(67). Circus was used to visualize genome map, transcripts, and expression levels. MHCs of *N. Communis* Nm2 were detected using the online MOTIF search tool of GenomeNet (https://www.genome.jp/tools/motif/MOTIF2.html).

### Construction of Sequence Datasets

The NCBI non-redundant protein database was queried using BLASTp and HMMER v3.3.2 for homologs of the MtrA, MtrB, MtrC, OMC, and PIO proteins in members of ammonia-oxidizing guilds having a sequenced genome. Homologs of MtrA, MtrB, and MtrC, were additionally queried in the NCBI protein nr database in prokaryotes having sequenced genomes. No MTR proteins were found in archaeal ammonia oxidizers. Sequences of each protein family were aligned using the MAFFT algorithm version 7.313 (https://mafft.cbrc.jp/alignment/software/) with parameters --ep 0 –genafpair --maxiterate 1000, and inspected manually for the presence of gaps and misalignment. Individual gene trees were constructed with IQ-Tree version 1.6.6, with the best fitting model chosen according to the BIC criterion with 1000 ultrafast bootstraps.

### Construction of Species Tree

A species tree based on single-copy universal genes was constructed for the 208 microbial species using the “bcgTree” (https://github.com/molbiodiv/bcgTree) pipeline, specifically including groups known to be ammonia oxidizers. Predicted protein-coding sequences of 208 species were quired against a database of hidden markov model (HMM) profiles of the single-copy universal genes (48; Supp. Table 6) found in over 95% of the proteomes using HMMER v3.3.2. Best matches remaining after a gene-specific cut-off (48; Supp. Table 6) were retained and aligned using MUSCLE v3.8.1551) (https://www.drive5.com/muscle/) with default settings. The resulting alignments were filtered with Gblocks v0.91b (http://molevol.cmima.csic.es/castresana/Gblocks.html) and concatenated with AMAS. (https://github.com/marekborowiec/AMAS). A maximum likelihood phylogenetic tree was constructed from the alignment with RAxML v8.2.12 (https://github.com/stamatak/standard-RAxML). A subtree of the constructed species tree was used in Figure 1.

### Construction of Gene Tree

The MtrA proteins retrieved from the proteomes in the species tree using HMMER v3.3.2, aligned using the MAFFT algorithm version 7.313. Individual MtrA tree was constructed with (i) IQ-Tree version 1.6.6 using the WAG as the chosen model (69) suitable for phylogenetic reconciliation with the best fitting model chosen according to the BIC criterion with 1000 ultrafast bootstraps and (ii) Phylobayes v4.1c using the WAG as the chosen model (from IQ-Tree) and using two chains.

### Divergence Time Estimation

Divergence time analyses were performed using Phylobayes v4.1c with a fixed topology from the RaxML composite alignment under the CAT20 model, birth–death and uniform tree priors, and uncorrelated gamma distributed (UGM), autocorrelated lognormal (LN) and autocorrelated CIR process rate models for both prior and posterior estimates. After chain convergence (effective size > 50, variable discrepancies < 0.30), the initial 20% of sampled generations were discarded as burn-in, and trees and posterior probability support values were calculated from completed chains. The sampled age estimates calculated with and without posterior likelihood calculations were compared, as suggested previously (58). A molecular clock was also applied to MtrA tree to support the reconciled root of the MtrA metabolism as described above using autocorrelated CIR process rate model.

### Age Constraints

Secondary calibrations were applied to the divergence times of archaeal and bacterial groups within the species tree and gene tree. For the species tree, we applied a time constraint for archaea within (i) *Methanosarcinales* derived from *Methanosarcinale* group and (ii) the split of *Methanococcoides* and *Methanohalobium* groups using the divergence time estimates based on the ribosomal alignment partitions by Wolfe et al. (42). We also applied time constraints for bacteria for the crown diversifications of (i) the genus *Aeromonas* (43) and (ii) the genus of *Vibrio* (44), and divergence of total-group *Nostocales* in our tree defined as the clade including *Nostoc, Fischerella*, and *Rivularia* (48). The root was calibrated with a prior of 3.9 ± 0.23 Ga ranging from 4.36–3.44 G as calculated previously (42). All calibrations are listed in Supplementary Table 7. To date, the MtrA trees, estimated ages of nine crown groups in the reconciled species-gene tree, were used as secondary calibrations (Supp. Table 8).

### Phylogenetic Reconciliation

Phylogenetic reconciliation infers reticulate events of gene evolutionary history by comparing the gene tree to a species trees under evolutionary models accounting for events such as duplications, transfers, losses, and incomplete lineage sorting. (70). We applied ecceTERA v1.24 to reconstruct species-tree-aware gene trees via the joint amalgamation method (55). The phylogenetic reconciliation was performed using the following sets of DTL (δ, τ, λ) event cost vectors: (1, 1, 1) = A, (1, 3, 1) = B, without (A00, B00) and with (A10, B10) transfers to/from a dead lineage, or the Pareto-optimal strategies 1 = C01 and 3 = C03, as previously published (55, 71, 72). A reconciliation viewer, SylvX (http://www.sylvx.org/), was used methods to ease the interpretation and comparison of reconciliations.

### Application of horizontal gene transfer-based evolutionary model assessment

HGT events to key nodes were selected using the MtrA gene and species tree reconciliations (Supp. Fig.5). For each HGT selected, ’compatible’ chronograms sampled after burn-in showing the donor being older than the recipient were selected. The percentage of compatible chronograms was then calculated for each HGT (Supp. Table 5) as described by Fournier et al. (48). Posterior age distributions were then generated from selected compatible chronograms.

## ACKNOWLEDGMENTS

We are thankful for the use of the HARVARD CNS electron microscopy facility. We thank Nicki Watson for her effort on TEM and Edward LaBelle for electrochemical reactor design. We thank Lisa Stein for providing *Nitrosomonas Communis* Nm2 strain. We also thank Jennifer Delaney for logistical support and Jo Wolfe for valuable discussion. This work and AG were financially supported by the European Union Marie Sklodowska-Curie individual global fellowship (MSCA-IF-GF) PAERADOX, Grant no: 800364.

## DATA DEPOSITION

Supplementary data files are available at https://github.com/ardagulay/.

## REFERENCES

1. M. Dworkin, S. Falkow, R. Eugene, S. Karl-Heinz, E. Stackebrandt, The Prokaryotes, M. Dworkin, S. Falkow, E. Rosenberg, K.-H. Schleifer, E. Stackebrandt, Eds. (Springer New York, 2006).

2. H. Koch, et al., Expanded metabolic versatility of ubiquitous nitrite-oxidizing bacteria from the genus Nitrospira. Proc. Natl. Acad. Sci. 112, 11371–11376 (2015).

3. B. Bayer, et al., Metabolic versatility of the nitrite-oxidizing bacterium Nitrospira marina and its proteomic response to oxygen-limited conditions. ISME J. 15, 1025–1039 (2021).

4. M. Berney, C. Greening, R. Conrad, W. R. Jacobs, G. M. Cook, An obligately aerobic soil bacterium activates fermentative hydrogen production to survive reductive stress during hypoxia. Proc. Natl. Acad. Sci. 111, 11479–11484 (2014).

5. L. Shi, et al., Extracellular electron transfer mechanisms between microorganisms and minerals. Nat. Rev. Microbiol. (2016) https://doi.org/10.1038/nrmicro.2016.93.

6. S. E. McGlynn, G. L. Chadwick, C. P. Kempes, V. J. Orphan, Single cell activity reveals direct electron transfer in methanotrophic consortia. Nature 526, 531–535 (2015).

7. X. Deng, N. Dohmae, K. H. Nealson, K. Hashimoto, A. Okamoto, Multi-heme cytochromes provide a pathway for survival in energy-limited environments. Sci. Adv. 4, 1–9 (2018).

8. D. R. Lovley, J. D. Coates, E. L. Blunt-Harris, E. J. P. Phillips, J. C. Woodward, Humic substances as electron acceptors for microbial respiration. Nature 382, 445–448 (1996).

9. D. R. Lovley, E. J. P. Phillips, Y. A. Gorby, E. R. Landa, Microbial reduction of uranium. Nature 350, 413–416 (1991).

10. J. A. Gralnick, H. Vali, D. P. Lies, D. K. Newman, Extracellular respiration of dimethyl sulfoxide by Shewanella oneidensis strain MR-1. Proc. Natl. Acad. Sci. U. S. A. 103, 4669–4674 (2006).

11. D. R. Bond, D. E. Holmes, L. M. Tender, D. R. Lovley, Electrode-reducing microorganisms that harvest energy from marine sediments. Science (80-.). 295, 483–485 (2002).

12. J. Raymond, The Effect of Oxygen on Biochemical Networks and the Evolution of Complex Life. Science (80-.). 311, 1764–1767 (2006).

13. L. A. David, E. J. Alm, Rapid evolutionary innovation during an Archaean genetic expansion. Nature 469, 93–96 (2011).

14. H. Koops, U. Purkhold, A. Pommerening-Röser, G. Timmermann, M. Wagner, “The Lithoautotrophic Ammonia-Oxidizing Bacteria” in The Prokaryotes, 2nd Ed., M. Dworkin, S. Falkow, E. Rosenberg, K.-H. Schleifer, E. Stackebrandt, Eds. (Springer New York, 2006), pp. 778– 811.

15. E. Garcia-Robledo, et al., Cryptic oxygen cycling in anoxic marine zones. Proc. Natl. Acad. Sci. U. S. A. 114, 8319–8324 (2017).

16. T. E. Freitag, J. I. Prosser, Community Structure of Ammonia-Oxidizing Bacteria within Anoxic Marine Sediments. Appl. Environ. Microbiol. 69, 1359–1371 (2003).

17. R. Mortimer, et al., Anoxic nitrification in marine sediments. Mar. Ecol. Prog. Ser. 276, 37–52 (2004).

18. R. Yu, K. Chandran, Strategies of Nitrosomonas europaea 19718 to counter low dissolved oxygen and high nitrite concentrations. BMC Microbiol. 10, 70 (2010).

19. J. Geets, N. Boon, W. Verstraete, Strategies of aerobic ammonia-oxidizing bacteria for coping with nutrient and oxygen fluctuations. FEMS Microbiol. Ecol. 58, 1–13 (2006).

20. R. Khadka, et al., Evolutionary History of Copper Membrane Monooxygenases. Front. Microbiol. 9, 1–13 (2018).

21. Y. Wang, et al., A methylotrophic origin of methanogenesis and early divergence of anaerobic multicarbon alkane metabolism. Sci. Adv. 7 (2021).

22. Y. Wang, et al., The late Archaean to early Proterozoic origin and evolution of anaerobic methane-oxidizing archaea. mLife 1, 96–100 (2022).

23. I. Schmidt, R. J. M. van Spanning, M. S. M. Jetten, Denitrification and ammonia oxidation by Nitrosomonas europaea wild-type, and NirK-and NorB-deficient mutants. Microbiology 150, 4107– 4114 (2004).

24. L. J. Shaw, et al., Nitrosospira spp. can produce nitrous oxide via a nitrifier denitrification pathway. Environ. Microbiol. 8, 214–222 (2006).

25. E. Bock, I. Schmidt, R. Striven, D. Zart, Nitrogen loss caused by denitrifying Nitrosomonas cells using ammonium or hydrogen as electron donors and nitrite as electron acceptor. Arch Microbiol. 163, 16–20 (1995).

26. J. J. L. Cantera, L. Y. Stein, Role of nitrite reductase in the ammonia-oxidizing pathway of Nitrosomonas europaea. Arch. Microbiol. 188, 349–54 (2007).

27. L. Y. Stein, “Heterotrophic Nitrification and Nitrifier Denitrification” in Nitrification, (American Society of Microbiology, 2011), pp. 95–114.

28. A. Vilajeliu-Pons, et al., Microbial electricity driven anoxic ammonium removal. Water Res. 130, 168–175 (2018).

29. A. Kumar, et al., The ins and outs of microorganism-electrode electron transfer reactions. Nat. Rev. Chem. 1, 1–13 (2017).

30. D. S. Gruen, J. M. Wolfe, G. P. Fournier, Paleozoic diversification of terrestrial chitin-degrading bacterial lineages. BMC Evol. Biol. 19, 1–19 (2019).

31. D. Coursolle, J. A. Gralnick, Modularity of the Mtr respiratory pathway of Shewanella oneidensis strain MR-1. Mol. Microbiol. 77, 995–1008 (2010).

32. G. A. Coleman, et al., A rooted phylogeny resolves early bacterial evolution. Science (80-.). 372 (2021).

33. C. E. Turick, S. Shimpalee, P. Satjaritanun, J. Weidner, S. Greenway, Convenient non-invasive electrochemical techniques to monitor microbial processes: current state and perspectives. Appl. Microbiol. Biotechnol. 103, 8327–8338 (2019).

34. J. A. Kozlowski, J. Price, L. Y. Stein, Revision of N 2 O-Producing Pathways in the Ammonia-Oxidizing Bacterium Nitrosomonas europaea ATCC 19718. Appl. Environ. Microbiol. 80, 4930– 4935 (2014).

35. S. Kato, K. Hashimoto, K. Watanabe, Microbial interspecies electron transfer via electric currents through conductive minerals. Proc. Natl. Acad. Sci. 109, 10042–10046 (2012).

36. H. Richter, et al., Cyclic voltammetry of biofilms of wild type and mutant Geobacter sulfurreducens on fuel cell anodes indicates possible roles of OmcB, OmcZ, type IV pili, and protons in extracellular electron transfer. Energy Environ. Sci. 2, 506–516 (2009).

37. L. Chen, et al., Electron Communication of Bacillus subtilis in Harsh Environments. iScience 12, 260–269 (2019).

38. M. J. Marshall, et al., c-Type Cytochrome-Dependent Formation of U(IV) Nanoparticles by Shewanella oneidensis. PLoS Biol. 4, e268 (2006).

39. R. S. Hartshorne, et al., Characterization of an electron conduit between bacteria and the extracellular environment. Proc. Natl. Acad. Sci. U. S. A. 106, 22169–22174 (2009).

40. J. D. Caranto, A. C. Vilbert, K. M. Lancaster, Nitrosomonas europaea cytochrome P460 is a direct link between nitrification and nitrous oxide emission. Proc. Natl. Acad. Sci. 113, 14704–14709 (2016).

41. C. Magnabosco, K. R. Moore, J. M. Wolfe, G. P. Fournier, Dating phototrophic microbial lineages with reticulate gene histories. Geobiology 16, 179–189 (2018).

42. J. M. Wolfe, G. P. Fournier, Horizontal gene transfer constrains the timing of methanogen evolution. Nat. Ecol. Evol. 2, 897–903 (2018).

43. A. Sanglas, V. Albarral, M. Farfán, J. G. Lorén, M. C. Fusté, Evolutionary roots and diversification of the genus Aeromonas. Front. Microbiol. 8, 1–13 (2017).

44. H. Lin, M. Yu, X. Wang, X. H. Zhang, Comparative genomic analysis reveals the evolution and environmental adaptation strategies of vibrios. BMC Genomics 19, 1–14 (2018).

45. R. J. Horodyski, J. Allan Donaldson, Microfossils from the Middle Proterozoic Dismal Lakes Groups, Arctic Canada. Precambrian Res. 11, 125–159 (1980).

46. K. R. Moore, et al., An Expanded Ribosomal Phylogeny of Cyanobacteria Supports a Deep Placement of Plastids. Front. Microbiol. 10, 1–14 (2019).

47. M. Dos Reis, et al., Uncertainty in the Timing of Origin of Animals and the Limits of Precision in Molecular Timescales. Curr. Biol. 25, 2939–2950 (2015).

48. G. P. Fournier, et al., The Archean origin of oxygenic photosynthesis and extant cyanobacterial lineages. Proc. R. Soc. B Biol. Sci. 288, 1–10 (2021).

49. N. J. Planavsky, et al., Low Mid-Proterozoic atmospheric oxygen levels and the delayed rise of animals. Science (80-.). 346, 635–638 (2014).

50. H. D. Holland, The oxygenation of the atmosphere and oceans. Philos. Trans. R. Soc. B Biol. Sci. 361, 903–915 (2006).

51. C. T. Reinhard, N. J. Planavsky, The History of Ocean Oxygenation. Ann. Rev. Mar. Sci. 14, 331– 353 (2022).

52. M. Ren, et al., phylogenomics suggests oxygen availability as a driving force in Thaumarchaeota evolution. ISME J. 13, 2150–2161 (2019).

53. M. Stolzer, K. Siewert, H. Lai, M. Xu, D. Durand, Event inference in multidomain families with phylogenetic reconciliation. BMC Bioinformatics 16, 1–20 (2015).

54. R. Libeskind-Hadas, Y.-C. Wu, M. S. Bansal, M. Kellis, Pareto-optimal phylogenetic tree reconciliation. Bioinformatics 30, i87–i95 (2014).

55. E. Jacox, C. Chauve, G. J. Szöllősi, Y. Ponty, C. Scornavacca, ecceTERA: comprehensive gene tree-species tree reconciliation using parsimony: Table 1. Bioinformatics 32, 2056–2058 (2016).

56. P. Puigbò, A. E. Lobkovsky, D. M. Kristensen, Y. I. Wolf, E. V. Koonin, Genomes in turmoil: Quantification of genome dynamics in prokaryote supergenomes. BMC Med. 12, 1–19 (2014).

57. B. Snel, P. Bork, M. A. Huynen, Genomes in Flux: The Evolution of Archaeal and Proteobacterial Gene Content. Genome Res. 12, 17–25 (2002).

58. M. L. Coleman, S. W. Chisholm, Ecosystem-specific selection pressures revealed through comparative population genomics. Proc. Natl. Acad. Sci. 107, 18634–18639 (2010).

59. S. Koskiniemi, S. Sun, O. G. Berg, D. I. Andersson, Selection-Driven Gene Loss in Bacteria. PLoS Genet. 8, e1002787 (2012).

60. J. Chun, et al., Comparative genomics reveals mechanism for short-term and long-term clonal transitions in pandemic Vibrio cholerae. Proc. Natl. Acad. Sci. 106, 15442–15447 (2009).

61. N. A. Moran, Microbial Minimalism. Cell 108, 583–586 (2002).

62. J. O. Andersson, S. G. Andersson, Insights into the evolutionary process of genome degradation. Curr. Opin. Genet. Dev. 9, 664–671 (1999).

63. E. Marsili, J. B. Rollefson, D. B. Baron, R. M. Hozalski, D. R. Bond, Microbial biofilm voltammetry: Direct electrochemical characterization of catalytic electrode-attached biofilms. Appl. Environ. Microbiol. 74, 7329–7337 (2008).

64. S. Andrews, FastQC: A Quality Control Tool for High Throughput Sequence Data. http://www.bioinformatics.babraham.ac.uk/projects/fastqc/ (2010).

65. A. M. Bolger, M. Lohse, B. Usadel, Trimmomatic: A flexible trimmer for Illumina sequence data. Bioinformatics 30, 2114–2120 (2014).

66. D. Kim, J. M. Paggi, C. Park, C. Bennett, S. L. Salzberg, Graph-based genome alignment and genotyping with HISAT2 and HISAT-genotype. Nat. Biotechnol. 37, 907–915 (2019).

67. S. Kovaka, et al., Transcriptome assembly from long-read RNA-seq alignments with StringTie2. bioRxiv, 1–13 (2019).

68. C. L. Dupont, et al., Genomic insights to SAR86, an abundant and uncultivated marine bacterial lineage. ISME J. 6, 1186–1199 (2012).

69. B. Q. Minh, J. Trifinopoulos, D. Schrempf, H. A. Schmidt, IQ-TREE version 1.6.8: Tutorials and Manual Phylogenomic software by maximum likelihood. Manual/tutorial (2018).

70. G. J. Szllosi, E. Tannier, V. Daubin, B. Boussau, The inference of gene trees with species trees. Syst. Biol. 64, e42–e62 (2015).

71. N. Waglechner, A. G. McArthur, G. D. Wright, Phylogenetic reconciliation reveals the natural history of glycopeptide antibiotic biosynthesis and resistance. Nat. Microbiol. 4, 1862–1871 (2019).

72. R. Libeskind-Hadas, Y.-C. Wu, M. S. Bansal, M. Kellis, Pareto-optimal phylogenetic tree reconciliation. Bioinformatics 30, i87–i95 (2014).

