## Supplementary File 1 for "Ancient electrogenic survival metabolism of *β -proteobacterial* ammonia oxidizers for oxygen deficiency"

*DRAFT Version

**Supplementary Table 1:** Model EET proteins used in this study for homology search

| Protein Uniprot ID | Protein Name | Strain |
|---|---|---|
| Q8E8S0 | CymA | Shewanella oneidensis (strain MR-1) |
| Q8EG33 | MtrC | Shewanella oneidensis (strain MR-1) |
| Q8EG34 | MtrB | Shewanella oneidensis (strain MR-1) |
| Q8EG35 | MtrA | Shewanella oneidensis (strain MR-1) |
| Q749L1 | OmcC | Geobacter sulfurreducens (strain ATCC 51573 / DSM 12127 / PCA) |
| Q748W7 | OmcA | Geobacter sulfurreducens (strain ATCC 51573 / DSM 12127 / PCA) |
| Q749K5 | OmcB | Geobacter sulfurreducens (strain ATCC 51573 / DSM 12127 / PCA) |
| Q74G83 | ppcB | Geobacter sulfurreducens (strain ATCC 51573 / DSM 12127 / PCA) |
| G5EBD6 | ppcA | Geobacter sulfurreducens (strain ATCC 51573 / DSM 12127 / PCA) |
| Q74FJ0 | OmcE | Geobacter sulfurreducens (strain ATCC 51573 / DSM 12127 / PCA) |
| Q74A86 | OmcS | Geobacter sulfurreducens (strain ATCC 51573 / DSM 12127 / PCA) |
| Q74DC2 | OmpB | Geobacter sulfurreducens (strain ATCC 51573 / DSM 12127 / PCA) |
| Q749T5 | OmpC | Geobacter sulfurreducens (strain ATCC 51573 / DSM 12127 / PCA) |
| A1EBT2 | PioA | Rhodopseudomonas palustris (strain TIE-1) |
| A1EBT3 | PioB | Rhodopseudomonas palustris (strain TIE-1) |
| A1EBT4 | PioC | Rhodopseudomonas palustris (strain TIE-1) |

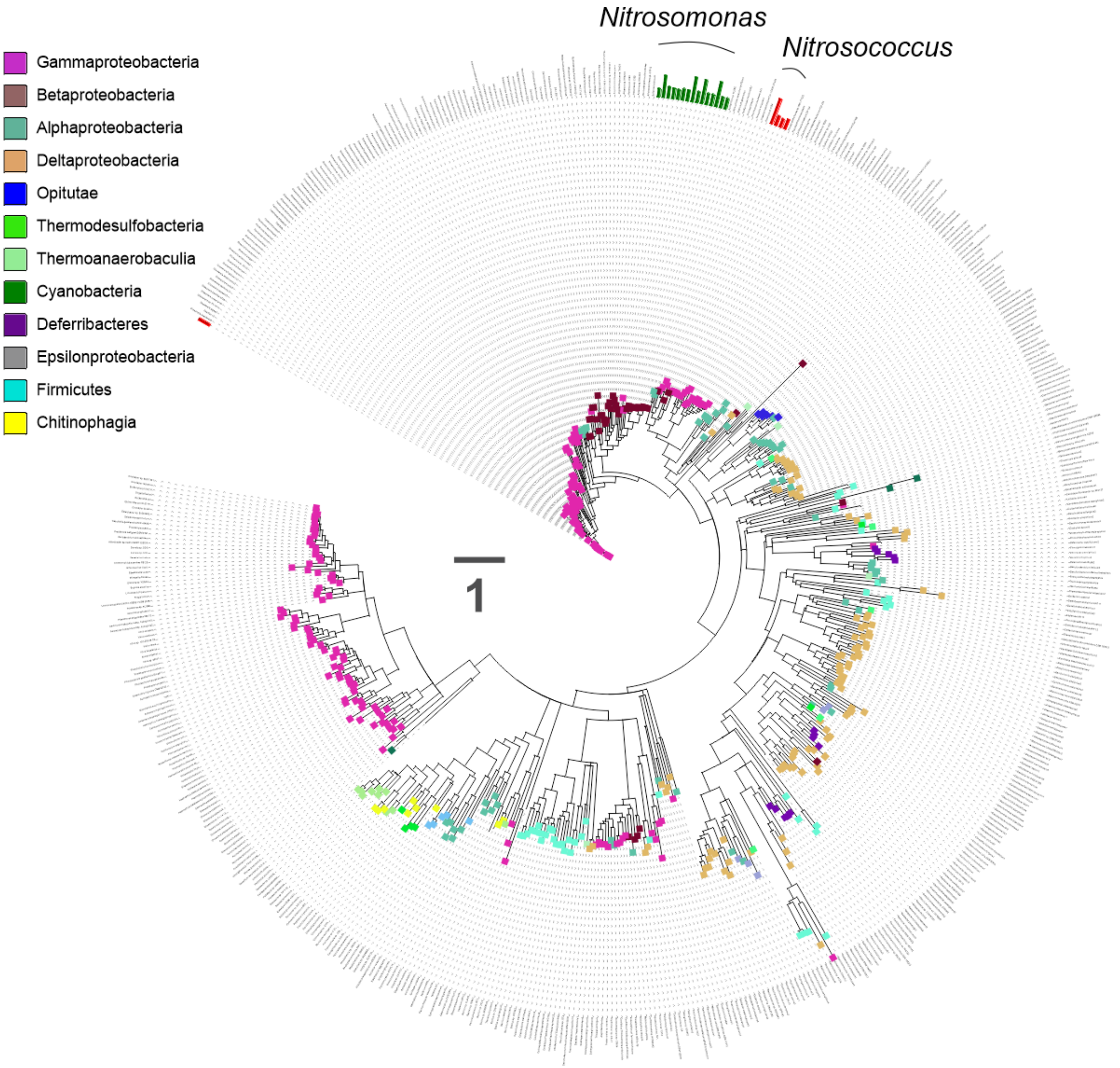


**Supplementary Figure 1**: The phylogenetic tree of homologs of periplasmic c-type cytochromes (mtrA) driving EET in *Shewanella Oneidensis* in the tree of bacterial life.

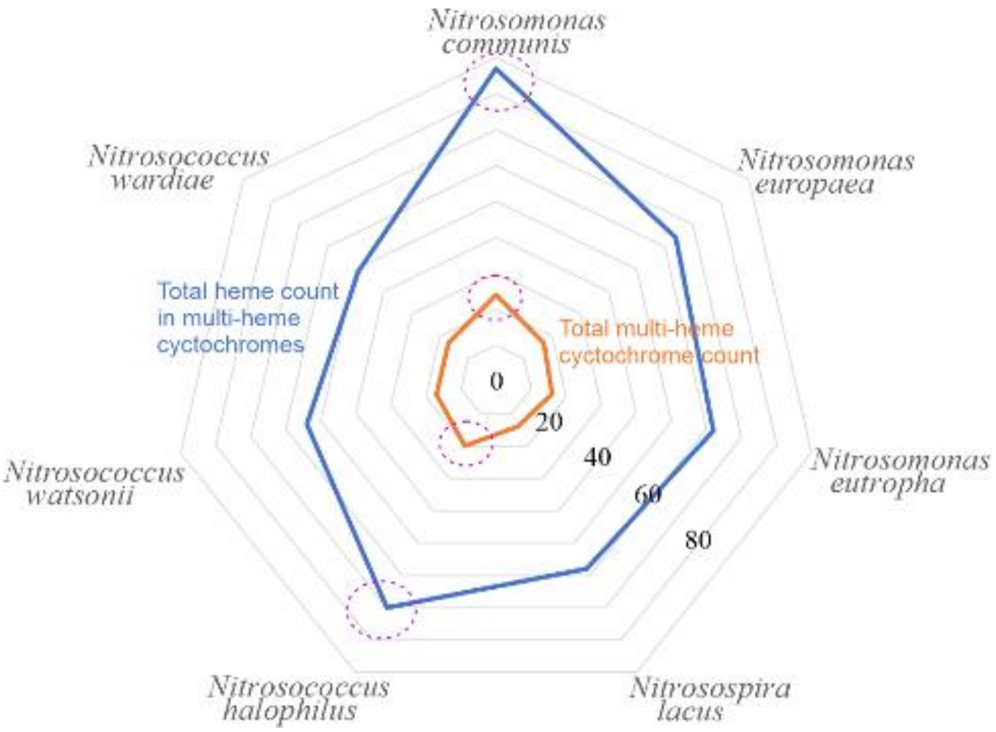


***Supplementary Figure 2*** *Comparison of total multi-heme cytochrome- and total heme-binding motif- counts of selected Nitrosomonas and Nitrosococcus strains*

**Supplementary Table 2:** Heme quantity and number of MHC in selected ammonia-oxidizing bacteria

| Nitrosomonas communis | Nitrosomonas europaea | Nitrosomonas eutropha | Nitrosospira lacus | Nitrosococcus halophilus | Nitrosococcus watsonii | Nitrosococcus wardiae |
|---|---|---|---|---|---|---|
| 10 | 8 | 8 | 8 | 10 | 8 | 8 |
| 8 | 8 | 8 | 8 | 10 | 8 | 5 |
| 8 | 8 | 8 | 8 | 8 | 5 | 4 |
| 8 | 4 | 4 | 4 | 5 | 4 | 4 |
| 4 | 4 | 4 | 4 | 4 | 4 | 3 |
| 4 | 4 | 4 | 4 | 4 | 3 | 3 |
| 4 | 4 | 4 | 4 | 3 | 2 | 2 |
| 4 | 4 | 4 | 4 | 2 | 2 | 2 |
| 4 | 3 | 3 | 3 | 2 | 2 | 2 |
| 3 | 2 | 2 | 2 | 2 | 2 | 2 |
| 3 | 2 | 2 | 2 | 2 | 2 | 2 |
| 3 | 2 | 2 | 2 | 2 | 2 | 2 |
| 2 | 3 | 2 | 2 | 2 | 2 | 2 |
| 2 | 2 | 2 | 3 | 2 | 2 | 2 |
| 2 | 2 | 3 | | 2 | 2 | 2 |
| 2 | 2 | 2 | | 2 | 2 | 2 |
| 2 | 2 | | | 2 | 2 | 2 |
| 2 | | | | 2 | | |
| 2 | | | | 2 | | |
| 2 | | | | 2 | | |
| 2 | | | | | | |
| 2 | | | | | | |
| 2 | | | | | | |
| 2 | | | | | | |

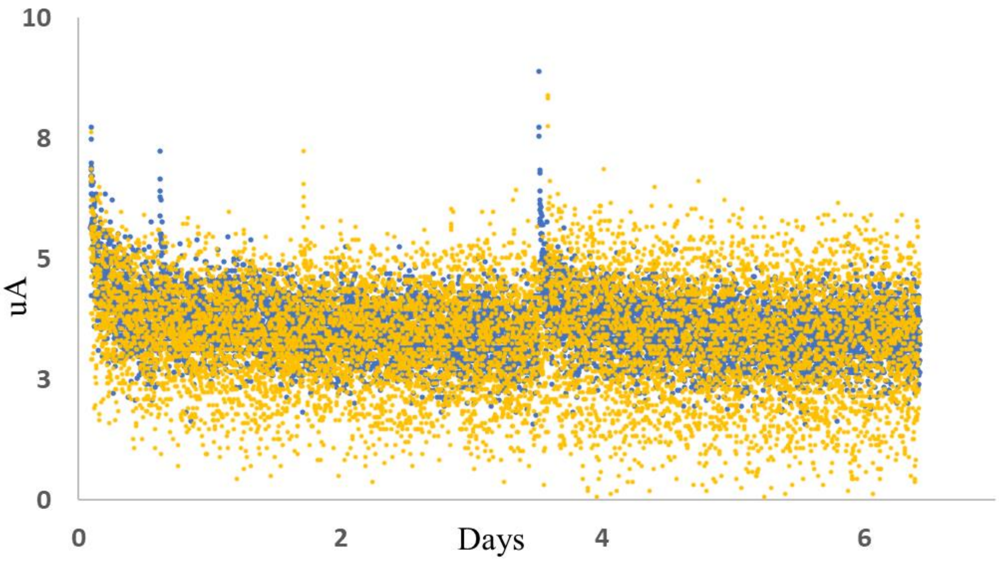


**Supplementary Figure 3**: Electrochemical characteristics of *Nitrosococcus Halophilus* on graphite felt electrode. Anodic currents were measured using an electrode poised at 0.3V versus AgCl supplemented with X mM $NH_4^+$. Control (yellow color) yielded similar current values as live incubations.

**Supplementary Table 3** Detected multi-heme cytochromes in *Nitrosomonas communis* Nm2

| Gene ID | Description | Genome positions of heme motifs (C-X-X-C-H) in the cyctochrome | Total Heme motif Count |
|---|---|---|---|
| AAW31_00475 | cytochrome C | 40..44,71..75,111..115,135..139,158..162,184..188,208..212,230..234,253..257,282..286 | 10 |
| AAW31_01285 | hydroxylamine reductase | 114..118,180..184,207..211,264..268,274..278,294..298,345..349,395..399 | 8 |
| AAW31_16290 | hydroxylamine reductase | 103..107,169..173,196..200,253..257,263..267,283..287,334..338,384..388 | 8 |
| AAW31_18275 | hydroxylamine reductase | 114..118,180..184,207..211,264..268,274..278,294..298,345..349,395..399 | 8 |
| AAW31_01270 | cytochrome C | 39..43,67..71,131..135,159..163 | 4 |
| AAW31_01275 | cytochrome C554 | 36..40,85..89,113..117,159..163 | 4 |
| AAW31_16275 | cytochrome C | 39..43,67..71,131..135,159..163 | 4 |
| AAW31_16280 | cytochrome C554 | 36..40,85..89,113..117,159..163 | 4 |
| AAW31_18265 | cytochrome C554 | 36..40,85..89,113..117,159..163 | 4 |
| AAW31_00995 | alcohol dehydrogenase | 34..38,180..184,306..310 | 3 |
| AAW31_10965 | alcohol dehydrogenase | 59..63,205..209,331..335 | 3 |
| AAW31_14635 | molecular chaperone DnaJ | 147..151,186..190,200..204 | 3 |
| AAW31_00285 | hypothetical protein | 53..57,290..294 | 2 |
| AAW31_00560 | cytochrome C | 126..130,333..337 | 2 |
| AAW31_01000 | cytochrome C553 | 37..41,139..143 | 2 |
| AAW31_05525 | cytochrome C | 63..67,207..211 | 2 |
| AAW31_08575 | alcohol dehydrogenase | 39..43,95..99 | 2 |
| AAW31_09880 | lipoprotein | 130..134,305..309 | 2 |
| AAW31_10270 | hypothetical protein | 113..117,351..355 | 2 |
| AAW31_10960 | cytochrome C553 | 17..21,119..123 | 2 |
| AAW31_11945 | cytochrome C | 32..36,136..140 | 2 |
| AAW31_12750 | cytochrome B6 | 298..302,428..432 | 2 |
| AAW31_14180 | hypothetical protein | 85..89,374..378 | 2 |
| AAW31_14525 | hypothetical protein | 249..253,494..498 | 2 |

**Supplementary Table 4:** Genes affiliated with anaerobic energy metabolism in *Nitrosomonas communis* Nm2 genome

| GENE_ID | DEFINITON | START | END | | FPKM |
|---|---|---|---|---|---|
| AAW31_00880 | cytochrome P460 | 205957 | 206562 | reverse | 323.39 |
| AAW31_02040 | cytochrome P460 | 472007 | 472603 | forward | 0.00 |
| AAW31_00985 | K00392 sulfite reductase (ferredoxin) [EC:1.8.7.1] \| (GenBank) nitrite reductase | 229655 | 231751 | reverse | 33.50 |
| AAW31_10555 | K02305 nitric oxide reductase subunit C \| (GenBank) cytochrome C | 2330479 | 2330931 | forward | 0.00 |
| AAW31_10560 | K04561 nitric oxide reductase subunit B [EC:1.7.2.5] \| (GenBank) nitric oxide reductase | 2330964 | 2332310 | forward | 0.00 |
| AAW31_10565 | K04748 nitric oxide reductase NorQ protein \| (GenBank) ATPase AAA | 2332323 | 2333147 | forward | 0.00 |
| AAW31_13320 | K04748 nitric oxide reductase NorQ protein \| (GenBank) ATPase AAA | 2946222 | 2947025 | forward | 0.00 |
| AAW31_13325 | K02448 nitric oxide reductase NorD protein \| (GenBank) von Willebrand factor A | 2947069 | 2949396 | forward | 1.39 |
| AAW31_06015 | K07234 uncharacterized protein involved in response to NO \| (GenBank) NnrS family protein | 1322333 | 1323571 | reverse | 0.00 |
| AAW31_04320 | K07234 uncharacterized protein involved in response to NO \| (GenBank) NnrS family protein | 983156 | 984349 | forward | 0.00 |

Supplementary Table 5: Genes affiliated with carbon fixation metabolism in *Nitrosomonas communis Nm2* genome

| GENE_ID | DEFINITON | START | END | | FPKM |
|---|---|---|---|---|---|
| AAW31_03085 | K01783 ribulose-phosphate 3-epimerase [EC:5.1.3.1] | 205957 | 206562 | reverse | 71.68 |
| AAW31_06725 | K01601 ribulose-bisphosphate carboxylase large chain [EC:4.1.1.39] | 472007 | 472603 | forward | 56.70 |
| AAW31_06730 | K01602 ribulose-bisphosphate carboxylase small chain [EC:4.1.1.39] | 229655 | 231751 | reverse | 242.05 |
| AAW31_13310 | K01601 ribulose-bisphosphate carboxylase large chain [EC:4.1.1.39] | 2330479 | 2330931 | forward | 11.17 |
| AAW31_13315 | K01602 ribulose-bisphosphate carboxylase small chain [EC:4.1.1.39] | 2330964 | 2332310 | forward | 0.00 |

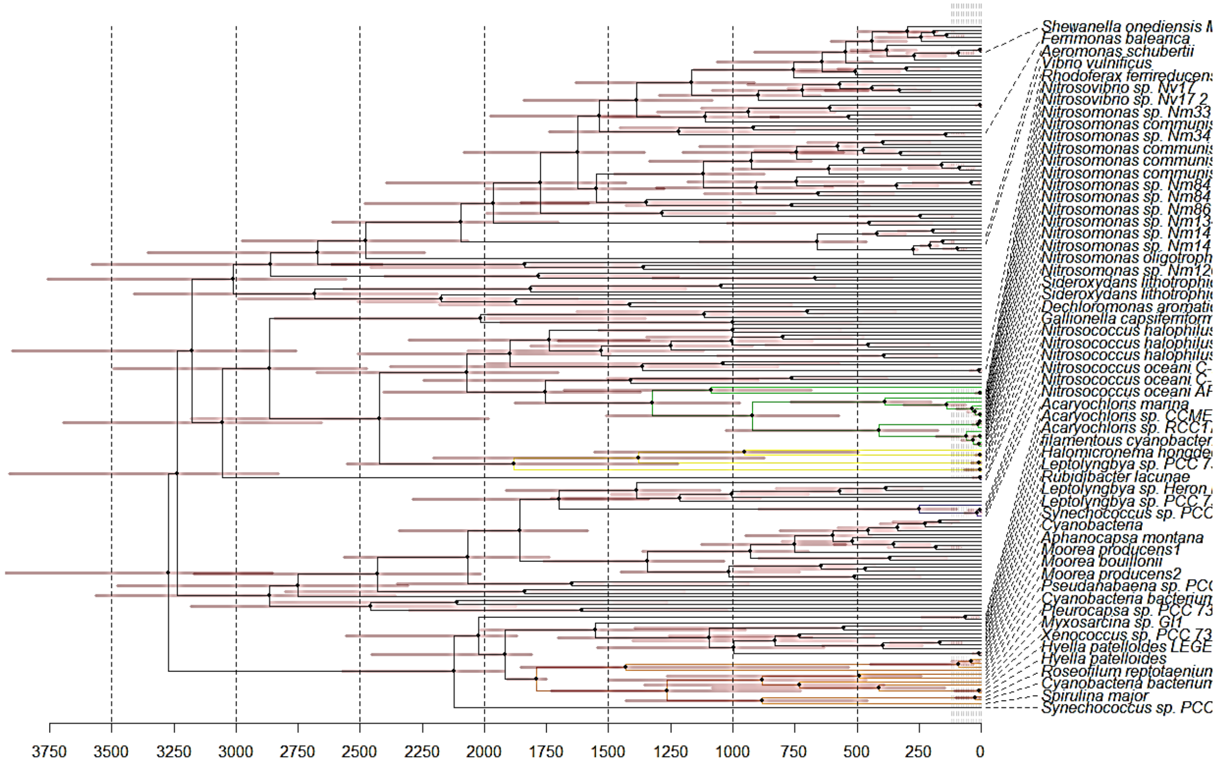


**Supplementary Figure 4** Chronogram depicting the phylogenetic relationships between mtrA homologs, and posterior age estimates obtained under the prior. Ages were estimated in PhyloBayes using the WAG substitution model and the UGM molecular clock model. Horizontal bars on the nodes indicate 95% CIs.

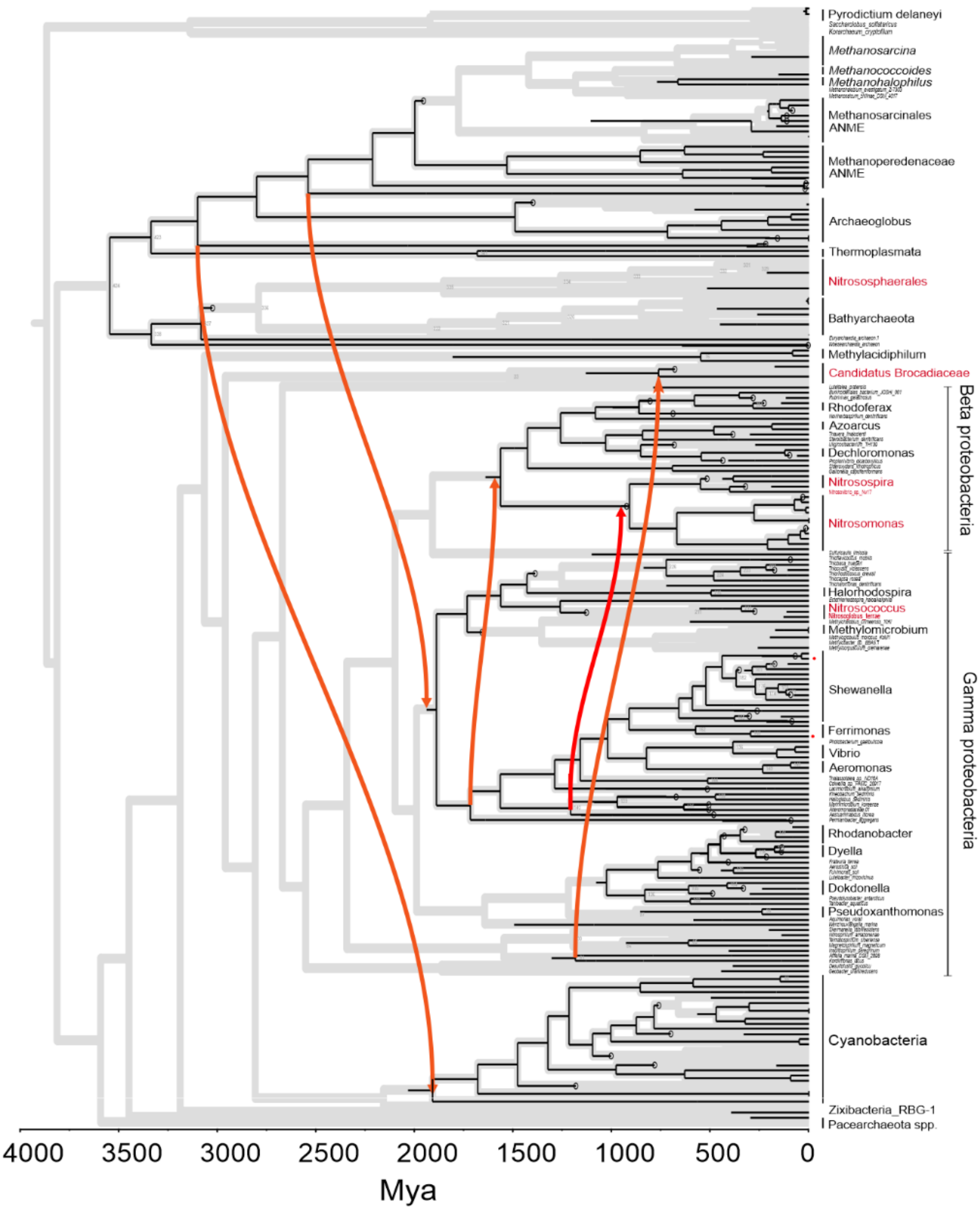


**Supplementary Figure 5** Map of index of selected HGTs on the bacterial species tree. HGT are mapped with arrows from older (donor) node to younger (recipient nodes).

**Supplementary Table 5:** Compatibility results based on the HGT events in Figure 1

| | Compatibility |
|---|---|
| CIR+BD | 0.93 |
| CIR+UNIFORM | 0.94 |
| LN+BD | 0.23 |
| LN+UNIFORM | 0.52 |
| UGAM+BD | 0.80 |
| wUGAM+UNIFORM | 0.72 |

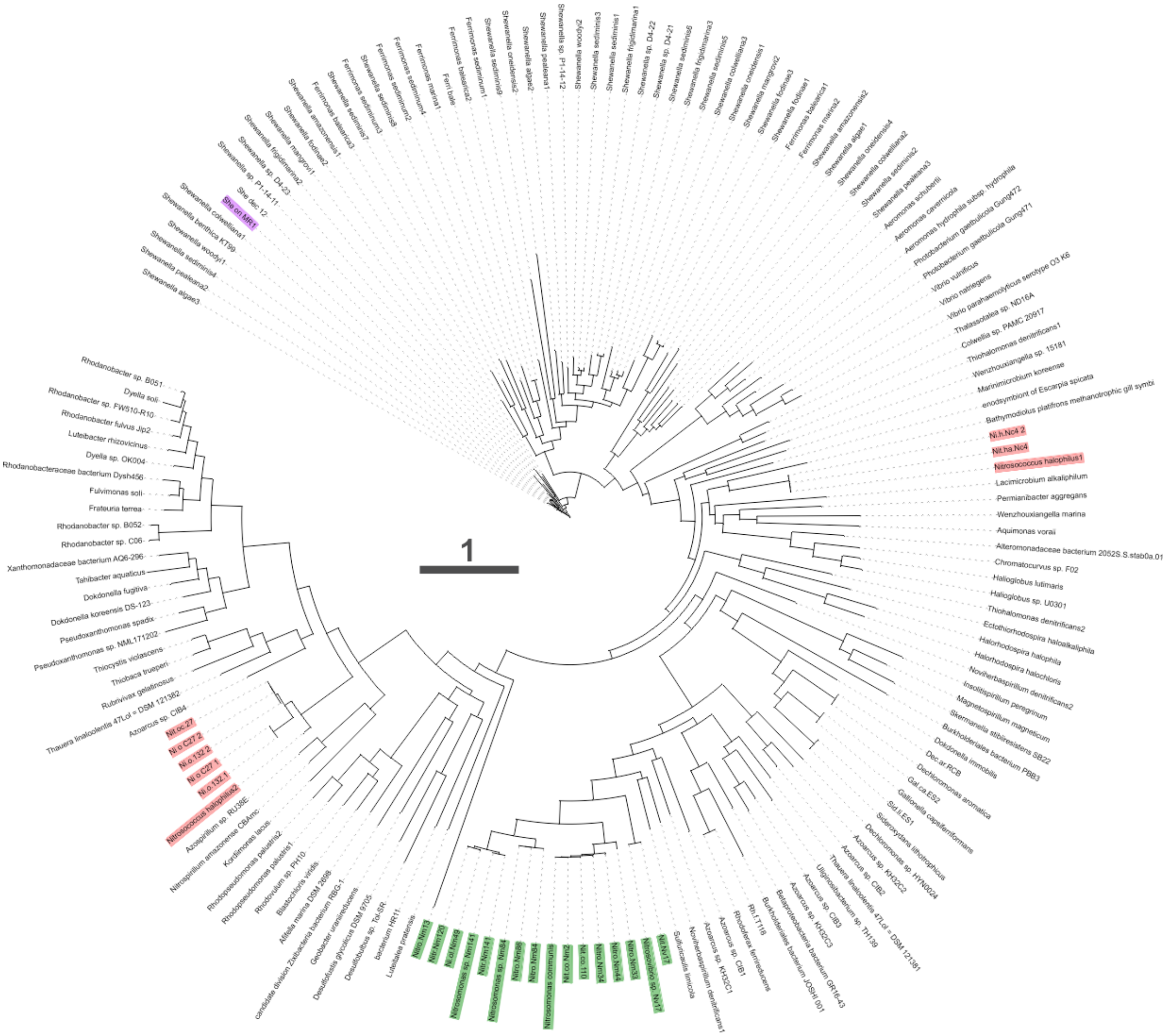


**Supplementary Figure 6:** The phylogenetic tree of homologs of integral outer-membrane β-barrel proteins (mtrB) driving EET in *Shewanella Oneidensis* in the tree of bacterial life.

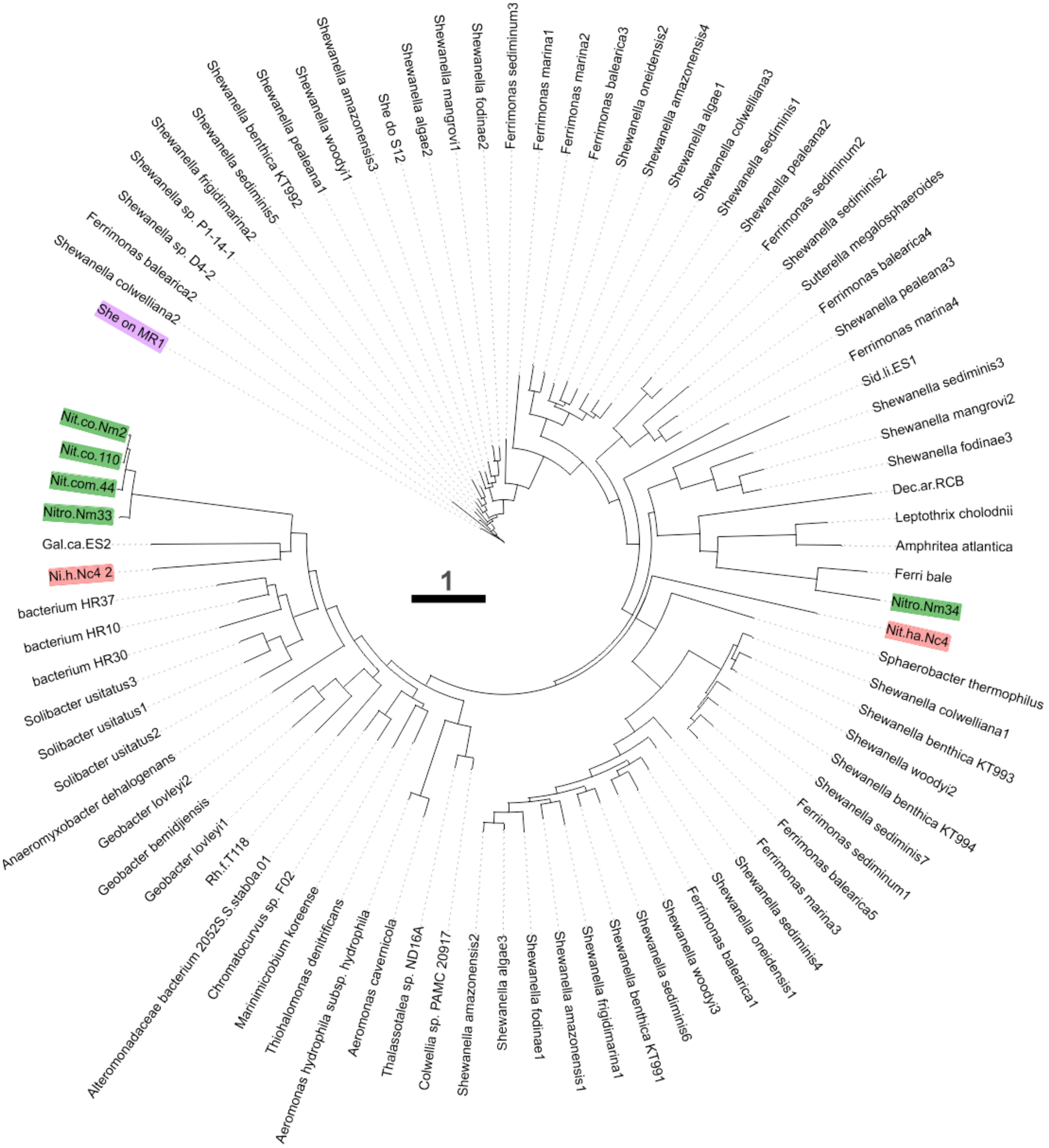


**Supplementary Figure 7:** The phylogenetic tree of homologs of outer-membrane-anchored c-type cytochromes (mtrC) driving EET in *Shewanella Oneidensis* in the tree of bacterial life.

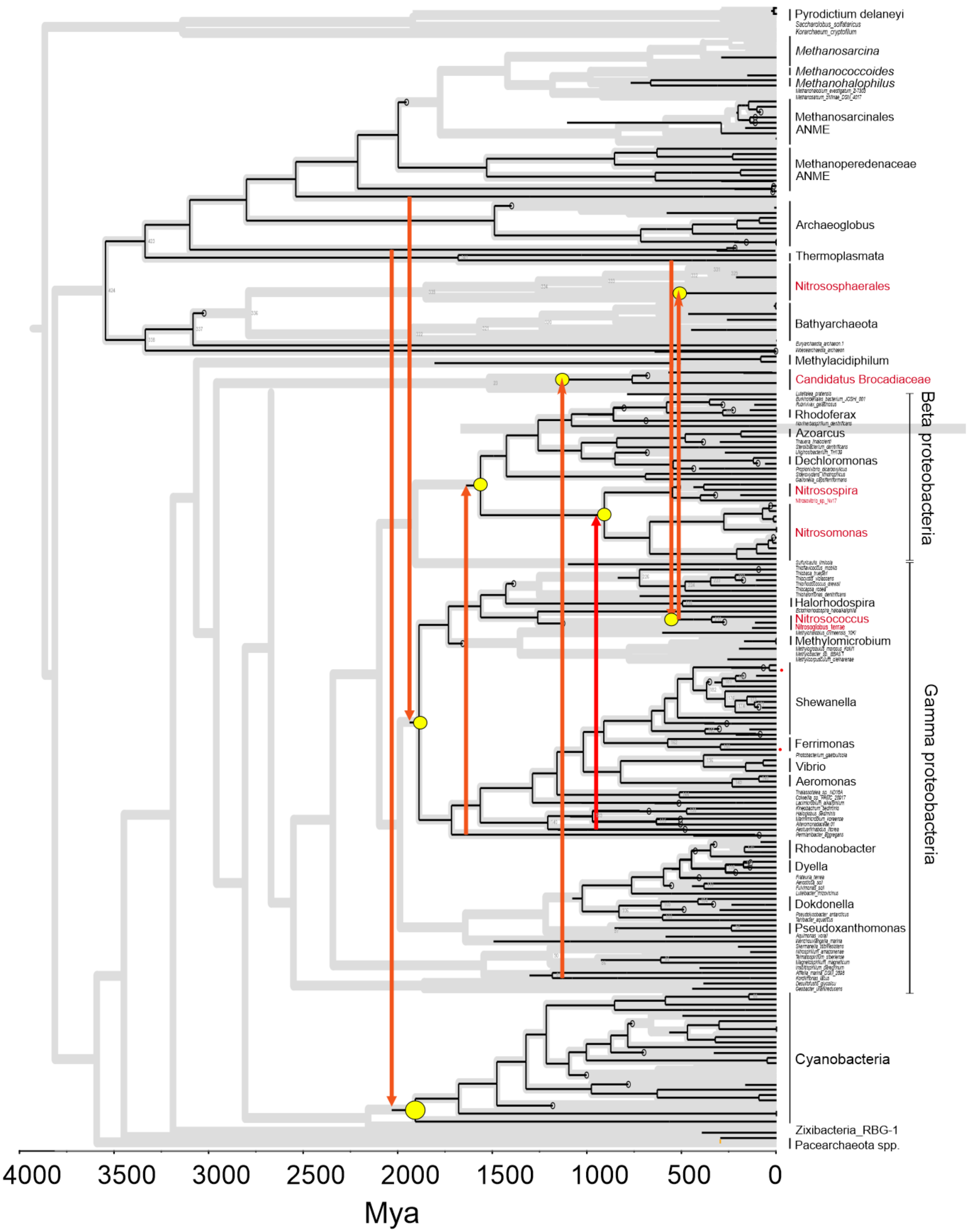


**Supplementary Figure 8** HGT receipent nodes

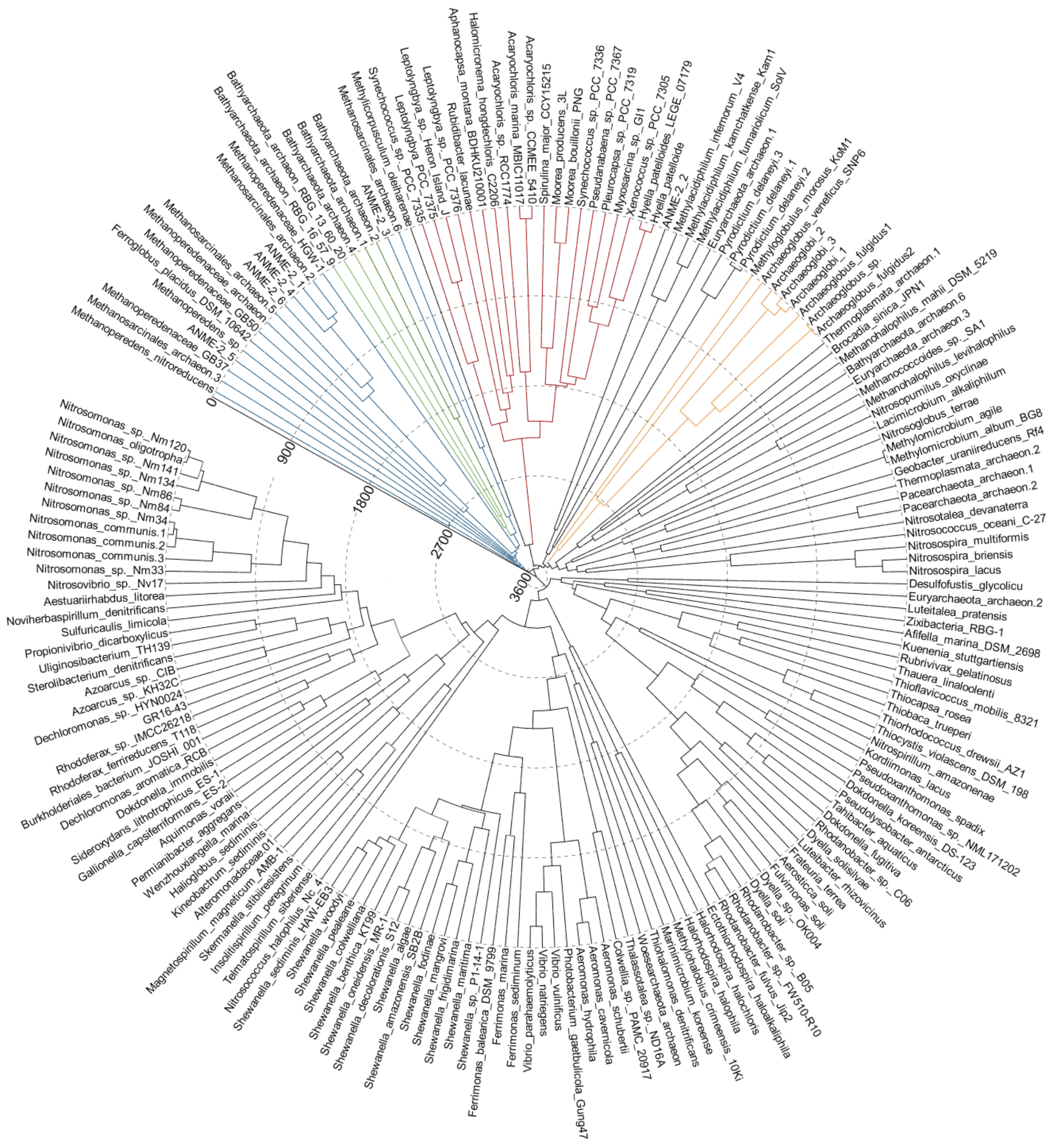


**Supplementary Figure 9** MtrA gene tree using the age estimates for the crown groups from the species tree as secondary calibrations

**Supplementary Table 6:** The single-copy genes searched in the proteomes of the species tree (48)

| Gene symbol | HMM | Cutoff of HMMER 2 | Cutoff for HMMER 3 | Description |
|---|---|---|---|---|
| alaS | TIGR00344 | 250 | 250 | Alanyl-tRNA synthetase |
| argS | PF00750 | -22.8 | 22.3 | Arginyl-tRNA synthetase |
| aspS | TIGR00459 | 265 | 600 | Aspartyl-tRNA synthetase |
| cgtA | TIGR02729 | 300 | 300 | Obg family GTPase CgtA |
| coaE | TIGR00152 | 37 | 37 | dephospho-CoA kinase |
| cysS | TIGR00435 | 250 | 250 | cysteinyl-tRNA synthetase |
| dnaA | TIGR00362 | 320 | 320 | chromosomal replication initiator protein DnaA |
| dnaG | TIGR01391 | 275 | 275 | DNA primase |
| dnaK | TIGR02350 | 1110 | 1110 | chaperone protein DnaK |
| dnaN | TIGR00663 | 30 | 30 | DNA polymerase III, beta subunit |
| dnaX | TIGR02397 | 200 | 150 | DNA polymerase III, subunits gamma and tau |
| engA | TIGR03594 | 250 | 250 | ribosome-associated GTPase EngA |
| era | TIGR00436 | 100 | 100 | GTP-binding protein Era |
| ffh | TIGR00959 | 540 | 540 | signal recognition particle protein |
| fmt | TIGR00460 | 160 | 160 | methionyl-tRNA formyltransferase |
| frr | TIGR00496 | 150 | 150 | ribosome recycling factor |
| ftsY | TIGR00064 | 290 | 290 | signal recognition particle-docking protein FtsY |
| glyS | TIGR00388 | 350 | 350 | glycyl-tRNA synthetase |
| glyS | TIGR00389 | 300 | 300 | glycyl-tRNA synthetase |
| gmk | TIGR03263 | 175 | 175 | guanylate kinase |
| grpE | PF01025 | -6.4 | 26.6 | co-chaperone GrpE |
| gyrA | TIGR01063 | 945 | 945 | DNA gyrase, A subunit |
| gyrB | TIGR01059 | 1170 | 1170 | DNA gyrase, B subunit |
| hisS | TIGR00442 | 270 | 270 | histidyl-tRNA synthetase |
| ileS | TIGR00392 | 600 | 600 | isoleucyl-tRNA synthetase |
| infB | TIGR00487 | 425 | 425 | translation initiation factor IF-2 |
| infC | TIGR00168 | 40 | 40 | translation initiation factor IF-3 |
| ksgA | TIGR00755 | 210 | 210 | dimethyladenosine transferase |
| lepA | TIGR01393 | 550 | 550 | GTP-binding protein LepA |
| leuS | TIGR00396 | 700 | 700 | leucyl-tRNA synthetase |
| ligA | TIGR00575 | 400 | 400 | DNA ligase, NAD-dependent |
| mnmA | TIGR00420 | 250 | 250 | tRNA (5-methylaminomethyl-2-thiouridylate)-methyltransf |
| mraW | PF01795 | -102.2 | 19.8 | MraW methylase family |
| nusA | TIGR01953 | 200 | 200 | transcription termination factor NusA |
| nusG | TIGR00922 | 140 | 140 | transcription termination/antitermination factor NusG |
| pgk | PF00162 | -15.7 | 21.9 | phosphoglycerate kinase |
| pheS | TIGR00468 | 280 | 280 | phenylalanyl-tRNA synthetase, alpha subunit |
| pheT | TIGR00471 | 220 | 220 | phenylalanyl-tRNA synthetase, beta subunit |
| pheT | TIGR00472 | 200 | 200 | phenylalanyl-tRNA synthetase, beta subunit |
| prfA | TIGR00019 | 475 | 475 | peptide chain release factor 1 |
| proS | TIGR00408 | 480 | 480 | prolyl-tRNA synthetase |
| proS | TIGR00409 | 300 | 300 | prolyl-tRNA synthetase |
| pyrG | TIGR00337 | 500 | 500 | CTP synthase |
| recA | TIGR02012 | 235 | 235 | recA protein |
| rfbA | TIGR00082 | 25 | 25 | ribosome-binding factor A |
| rnc | TIGR02191 | 165 | 165 | ribonuclease III |

**Supplementary Table 6:** Continue (48)

| | | | | |
|---|---|---|---|---|
| rplA | TIGR01169 | 200 | 200 | ribosomal protein L1 |
| rplB | TIGR01171 | 65 | 65 | ribosomal protein L2 |
| rplC | PF00297 | -79.6 | 26.8 | ribosomal protein L3 |
| rplD | PF00573 | -42.9 | 20.8 | ribosomal protein L4 |
| rplE | PF00281 | 6.2 | 20.9 | ribosomal protein L5 |
| rplF | PF00347 | 3.5 | 22.5 | ribosomal protein L6 |
| rplI | TIGR00158 | 50 | 50 | ribosomal protein L9 |
| rplJ | PF00466 | -12.8 | 21 | ribosomal protein L10 |
| rplK | TIGR01632 | 150 | 150 | ribosomal protein L11 |
| rplL | TIGR00855 | 65 | 65 | ribosomal protein L7/L12 |
| rplM | TIGR01066 | 30 | 30 | ribosomal protein L13 |
| rplN | TIGR01067 | 132 | 132 | ribosomal protein L14 |
| rplO | TIGR01071 | 60 | 60 | ribosomal protein L15 |
| rplP | TIGR01164 | 40 | 40 | ribosomal protein L16 |
| rplQ | TIGR00059 | 75 | 75 | ribosomal protein L17 |
| rplR | TIGR00060 | 80 | 80 | ribosomal protein L18 |
| rplS | TIGR01024 | 70 | 70 | ribosomal protein L19 |
| rplT | TIGR01032 | 19 | 19 | ribosomal protein L20 |
| rplU | TIGR00061 | 17 | 17 | ribosomal protein L21 |
| rplV | TIGR01044 | 60 | 60 | ribosomal protein L22 |
| rplW | PF00276 | 19.9 | 21.6 | ribosomal protein L23 |
| rplX | TIGR01079 | 16 | 16 | ribosomal protein L24 |
| rpmA | TIGR00062 | 110 | 110 | ribosomal protein L27 |
| rpmB | TIGR00009 | 45 | 45 | ribosomal protein L28 |
| rpmC | TIGR00012 | 8 | 8 | ribosomal protein L29 |
| rpmF | TIGR01031 | 13 | 13 | ribosomal protein L32 |
| rpmH | TIGR01030 | 30 | 30 | ribosomal protein L34 |
| rpmI | TIGR00001 | 40 | 40 | ribosomal protein L35 |
| rpoA | TIGR02027 | 180 | 180 | DNA-directed RNA polymerase, alpha subunit |
| rpoB | TIGR02013 | 1200 | 1200 | DNA-directed RNA polymerase, beta subunit |
| rpoC | TIGR02386 | 1250 | 1250 | DNA-directed RNA polymerase, beta' or beta'' subunit |
| rpoC | TIGR02387 | 1150 | 1150 | DNA-directed RNA polymerase, beta' or beta'' subunit |
| rpsB | TIGR01011 | -20 | -20 | ribosomal protein S2 |
| rpsC | TIGR01009 | 15 | 15 | ribosomal protein S3 |
| rpsD | TIGR01017 | 180 | 180 | ribosomal protein S4 |
| rpsE | TIGR01021 | 190 | 190 | ribosomal protein S5 |
| rpsF | TIGR00166 | 40 | 40 | ribosomal protein S6 |
| rpsG | TIGR01029 | 120 | 120 | ribosomal protein S7 |
| rpsH | PF00410 | -24.4 | 25.5 | ribosomal protein S8 |
| rpsI | PF00380 | -23.4 | 21.4 | ribosomal protein S9 |
| rpsJ | TIGR01049 | 70 | 70 | ribosomal protein S10 |
| rpsK | PF00411 | -29.8 | 21.8 | ribosomal protein S11 |
| rpsL | TIGR00981 | 80 | 80 | ribosomal protein S12 |
| rpsM | PF00416 | -5.6 | 21.3 | ribosomal protein S13 |
| rpsO | TIGR00952 | 30 | 30 | ribosomal protein S15 |
| rpsP | TIGR00002 | 35 | 35 | ribosomal protein S16 |
| rpsQ | PF00366 | 12.4 | 20.9 | ribosomal protein S17 |
| rpsR | TIGR00165 | 35 | 35 | ribosomal protein S18 |
| rpsS | TIGR01050 | 100 | 100 | ribosomal protein S19 |
| rpsT | TIGR00029 | 40 | 40 | ribosomal protein S20 |
| secA | TIGR00963 | 1000 | 1000 | preprotein translocase, SecA subunit |
| secE | TIGR00964 | 13 | 13 | preprotein translocase, SecE subunit |
| secG | TIGR00810 | 14 | 14 | preprotein translocase, SecG subunit |
| secY | TIGR00967 | 200 | 200 | preprotein translocase, SecY subunit |
| serS | TIGR00414 | 225 | 225 | seryl-tRNA synthetase |
| smpB | TIGR00086 | 110 | 100 | SmpB protein |
| thrS | TIGR00418 | 250 | 250 | threonyl-tRNA synthetase |
| tig | TIGR00115 | 75 | 75 | trigger factor |
| tilS | TIGR02432 | 80 | 80 | tRNA(Ile)-lysidine synthetase |
| tsf | TIGR00116 | 8 | 8 | translation elongation factor Ts |
| tyrS | TIGR00234 | 60 | 60 | tyrosyl-tRNA synthetase |
| uvrB | TIGR00631 | 1000 | 1000 | excinuclease ABC, B subunit |
| valS | TIGR00422 | 525 | 525 | valyl-tRNA synthetase |
| ybeY | TIGR00043 | 10 | 10 | conserved hypothetical protein YbeY |
| ychF | TIGR00092 | 225 | 225 | GTP-binding protein YchF |

**Supplementary Table 7:** Calibrations used in molecular clock models. All calibrations are listed in Ma. Taxon 1 and Taxon 2 refer to the taxa used in PhyloBayes commands

| **Node** | **Taxon I** | **Taxon II** | **Calibration** | **Reference** |
|---|---|---|---|---|
| Root | Korarchaeum cryptofilum | Shewanella sediminis | 4360-3440 | Fournier et al., 2021 |
| *Nostocales* Crown | Rivularia sp. PCC 7116 | Fischerella sp. PCC 9605 | 2070-1750 | Fournier et al., 2021 |
| *Methanococcoides* and *Methanohalobium* Split | Methanohalobium evestigatum | Methanosarcina burtonii | 1812-792 | Wolfe et al., 2018 |
| *Mehanosarcina* Crown | Methanosarcina barkeri | Methanosarcina mazei | 886-208 | Wolfe et al., 2018 |
| *Aeromonas* Crown | Aeromonas schubertii | Aeromonas cavernicola | 479-72 | Sanglas et al., 2017 |
| *Vibrio* Crown | Vibrio vulnificus | Vibrio natriegens | 278-113 | Lin et al., 2018 |

**Supplementary Table 8:** The single-copy genes searched in the proteomes of the species tree(48)

| CROWN GROUPS | Age range (both) | | Uni-CIR | | UNI-BD | |
|---|---|---|---|---|---|---|
| | Min | Max | Min | Max | Min | Max |
| MethanoperedenaceaeANME | 2041 | 3008 | 2041 | 2898 | 2321 | 3008 |
| Archaeoglobus | 2013 | 3051 | 2337 | 3051 | 2013 | 3029 |
| Nitrosomonadecea | 1371 | 1973 | 1467 | 1973 | 1793 | 1371 |
| Rhodocyclales | 1414 | 2019 | 2019 | 1546 | 1838 | 1414 |
| Chromatiaceae | 1482 | 2337 | 2337 | 1847 | 2186 | 1482 |
| Shewanella | 1249 | 1686 | 1686 | 1276 | 1543 | 1249 |
| Vibrio | 465 | 869 | 869 | 474 | 818 | 465 |
| Aeromonas | 251 | 476 | 476 | 251 | 476 | 269 |
| Cyanobacteria | 2247 | 2722 | 2722 | 2247 | 2698 | 2372 |
